# Local heterogeneity analysis of crystallographic and cryo-EM maps using shell-approximation

**DOI:** 10.1101/2023.04.11.536420

**Authors:** Vladimir Y. Lunin, Natalia L. Lunina, Alexandre G. Urzhumtsev

## Abstract

In X-ray crystallography and cryo-EM, experimental maps can be heterogeneous, showing different regions of the structure with different level of details. In this work we interpret the heterogeneity in terms of two parameters, assigned individually for each atom, combining the conventional parameter of atomic displacement with the resolution of the atomic image in the map. We propose a local real-space procedure to estimate the values of these heterogeneity parameters, assuming that a fragment of the density map and preliminary values of atomic coordinates are given. The procedure is based on the representation of the atomic image in an analytical form, as a function of the inhomogeneity parameters and atomic coordinates. In this article, we report the results of the tests both with simulated maps and maps derived from experimental data. For simulated heterogeneous maps containing regions with different resolutions, the method determines the local map resolution near the atomic centers and the values of the atomic displacement parameter with reasonable accuracy. For experimental maps, obtained as a Fourier synthesis of a given global resolution, estimated values of the local resolution are close to the global one, and the values of the estimated displacement parameters are close to the respective values in the refined model. Shown examples of the application of the proposed method to the experimental crystallographic and cryo-EM maps can be seen as a practical proof of method.

## 1. Introduction

Electron density distribution corresponding to a given atomic model is a key element in real-space fitting and refinement of macromolecular structures in X-ray crystallography (Rupp, 2009; Urzhumtsev and Lunin, 2019). For an appropriate quantitative comparison of a theoretical map (calculated from an atomic model) with the experimental one, the theoretical map calculated from the available atomic model should mimic the distortions of the experimental one. Considering that experimental errors and data imperfections including their incompleteness already have been treated in the best possible way, the two main sources of distortions of the experimental map at the late stages are the dynamic and static variation of the atomic positions and the limited resolution with which the atoms are represented by the map.

Both the scale of the positional variation (positional uncertainty) and the resolution of atomic images may change within the sample. Inside the macromolecular region, one may attribute them to each atom individually. In a recent series of papers (Urzhumtsev and Lunin, 2022a, 2022b; Urzhumtsev et al., 2022; Urzhumtseva et al., 2023), an efficient procedure has been proposed to calculate images of individual atoms adapted to distortions of the experimental map, provided the coordinates, atomic displacement parameter (ADP), and the atomic image resolution (AIR). In this article, we discuss an approach to solve the inverse problem, that, given a map and atomic coordinates, to find the local ADP and AIR values for each atom of the structure. The values found reflect the level of heterogeneity of the experimental map and can be used as initial values for further crystallographic refinement of these parameters (Urzhumtsev and Lunin, 2022b). The proposed approach is currently aimed to the range from medium to high resolution (1.5 – 3.5 Å), does not consider anisotropy of atomic images, the presence of deformation density (Bragg, 1920) and ignores errors in atomic positions (*e*.*g*., Cruickshank, 1999; Gurusaran et al., 2014; and references therein). The capability of the method to reproduce these distortion parameters and experimental maps allows addressing the following step, namely a common refinement of atomic positions and these parameters.

The above considerations are also applicable to the experimental maps other than electron density distribution, for example, to maps of the electrostatic scattering potential studied by cryo-electron microscopy (cryo-EM), nuclear density in studies of neutron diffraction, and other maps which can be represented by a sum of the contributions of individual atoms. Below, to be short, we will call all such maps as ‘density distribution’ maps.

In such methods, the samples contain a huge number of the molecules, in which the equivalent atoms may occur in different positions (static disorder). Furthermore, the position of any atom may vary in time due to thermal motion (dynamic disorder). Dynamic and static disorder blur the density peaks in the maps which show an average molecular image and are usually modeled by convolution of the theoretical atomic density with a Gaussian function, individual for each atom. This function defines the probability of the displacement of the atom from its mean position. Below we consider the isotropic case only, specifying the width of the Gaussian distribution by a single parameter *B* (atomic displacement parameter). This parameter is related to the mean square displacement |**u**| of the atom in three-dimensional space by *B* = (8*π*^2^⟨|**u**|^2^⟩/3). The convolution in real space with such function is equivalent to multiplication of the atomic scattering factor by exp[−*B*(sin *θ*/*γ*)^2^] where 2*θ* is the angle between the incident and reflected X-ray beams and *γ* is the wavelength. Previously, some methods to calculate atomic images in the maps of a given resolution knowing the ADP values, and eventually to refine these values, have been suggested by Diamond (1971), Lunin and Urzhumtsev (1984), Chapman (1995), Mooij *et al*. (2006), Chapman *et al*. (2013), DiMaio *et al*. (2015), Pintilie *et al*. (2020).

The term ‘resolution’ has different meaning in different fields of science and different experimental techniques. In macromolecular crystallography, the most commonly, the resolution of density map is defined in terms of the Fourier transform **F**(**s**) of the density distribution (*e*.*g*., Rupp, 2009). When working with a crystal, real or virtual, the Fourier transform is reduced to a discrete set of Fourier coefficients (structure factors), corresponding to the reciprocal space lattice. Initially, a resolution value *d* is attributed to each individual reflection, to the corresponding scattering vector **s**, and to the structure factor **F**(**s**). Numerically, it is defined as *d* = |**s**|^−1^ = *γ*/(2 sin *θ*). Then, this term is extended to the sets of reciprocal space vectors **s** that include all (or almost all) vectors with 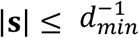. The value *d*_*min*_ is called the resolution of the respective data set which we note as *S*(*d*_*min*_). Similarly, this term *d*_*min*_ is applied to define the formal resolution of the map calculated as a Fourier synthesis

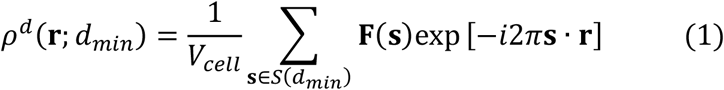

using such set *S*(*d*_*min*_) of reciprocal space lattice vectors. Here *V*_*cell*_ is the unit cell volume, **s · r** is the dot (scalar) product of two vectors. The resolution value *d* corresponding to a particular structure factor **F**(**s**) is numerically equal to the period of the associated Fourier harmonic *ψ*_**s**_(**r**) = exp[−*i*2*π***s · r**] in the direction **s**, so that the value *d*_*min*_ is sometimes referred to as the full-period Fourier resolution of the synthesis (1). A positional uncertainty, described by the convolution of *ρ*^*d*^(**r;** *d*_*min*_) with a Gaussian, is equivalent to a Gaussian weighting of structure factors reducing their magnitudes. To describe individual atomic uncertainties, such weight should be applied to individual atomic contributions included in **F**(**s**).

The truncation of the resolution limit causes blurring the peaks and appearance of Fourier series terminations ripples (James, 1948), a series of spherical shells of alternative sign map values, surrounding the atomic center. While not exceeding in amplitude, 10% of the height of the central peak, these ripples may produce visible distortions in maps in a large range of resolution, *e*.*g*., hiding density peaks for water molecules at resolution around 3 Å (Minichino et al., 2003) or producing false features mimicking deformation density in ultra-high-resolution studies (Afonine et al., 2004, Fig. 4). The effect of the truncation of resolution in Fourier synthesis (1) may be described as the convolution of the theoretical distribution with the three-dimensional interference function (Urzhumtsev and Lunin, 2022a).

In cryo-EM, a number of higher-resolution Fourier harmonics are either missed due to experimental features and image processing or negligibly small. The resolution here is usually estimated by comparing, in resolution shells, structure factors calculated from two maps (half-maps) reconstructed independently from two different subsets of experimental data (van Heel *et al*., 1982; Saxton and Baumeister, 1982; Rosenthal and Henderson, 2003; van Heel and Schatz, 2017). The key difference between the two definitions is that the crystallographic definition tells how many Fourier harmonics are present in the map, while the cryo-EM definitions based on the Fourier Shell Correlation function (FSC) tell how many of them are reliable.

There exist other approaches to define the resolution. Addressing the question of the data strength, the resolution may be declared as the limit beyond which excluding structure factors does not affect atomic model refinement (Karplus and Diederichs, 2012) or the maps themselves (Afonine *et al*., 2018). Alternatively, one may define the resolution value addressing directly to the map’s features. For example, one can analyzed the Model Trapping Function (Pintilie and Chiu, 2012; Lunin *et al*., 2019), which shows the number of atoms inside the region of the highest density values for variable cut-off values. Also, one can define the resolution from the ‘optical features’ of the map (Vaguine *et al*., 1999; Penczek, 2010; Urzhumtseva *et al*., 2013). Some discussions concerning comparison of different resolution definitions and respective terms can be found in Liao and Frank (2010), Penczek (2010), Evans and Murshudov (2013), Afonine *et al*. (2018). In what follows, we consider the term ‘resolution’ in the crystallographic meaning associated with (1).

The formal resolution value of Fourier synthesis (1) characterizes the macromolecular image in the map as a whole while details of different parts of this image may be different. For example, Fig. 1 shows an image of four tyrosine residues inside the same virtual crystal cell. This image was calculated as a single Fourier synthesis of a formal resolution equal to 2 Å. The fragment A reproduces the features of a typical 2 Å resolution map, while the fragment C looks like a 5 Å-resolution one. The fragment B demonstrates the effect of increasing ADP values of all atoms to 100 Å^2^. Fragment D reveals the features of images of a resolution higher than 2 Å, while actually being a sharpened (Terwilliger et al., 2018) 2 Å -resolution image. Thus, to evaluate properly the quality of a map, a single global characteristic is insufficient (Cardone et al., 2013) and some parameters characterizing local features are required (Chapman et al., 2013; Kucukelber et al., 2014; Vilas et al., 2018; Ramírez-Aportela et al., 2019).

**Fig. 1.**
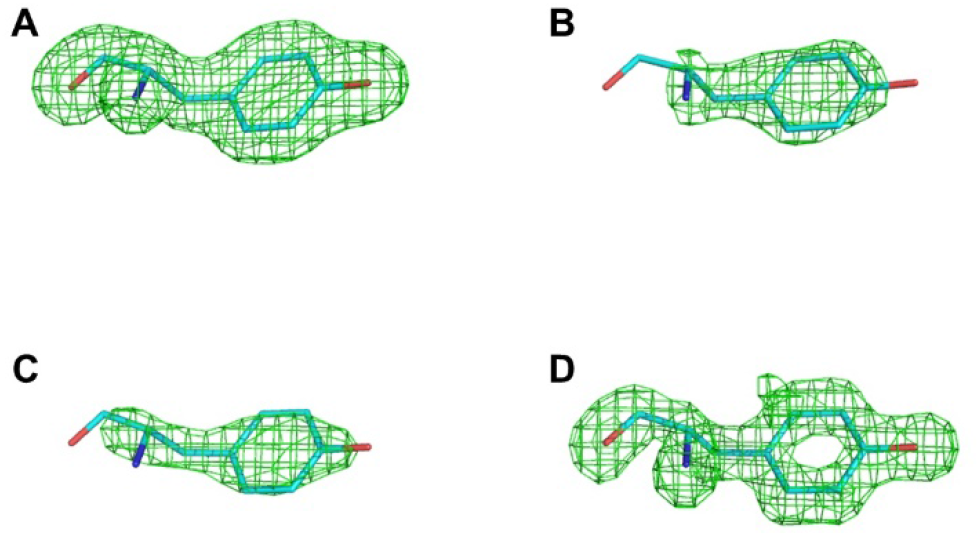
Fourier synthesis calculated as (1) with the resolution *d*_*min*_ = 2 Å reproduces the features of the nominal resolution map (A) and those of lower (B, C) and higher (D) resolution. Structure factors used to calculate this synthesis were the sums of structure factors from four sets, corresponding to the fragments: A) 2 Å-resolution set of structure factors for atoms with *B* = 20 Å^2^; B) 2 Å-resolution set of structure factors for atoms with *B* increased to 100 Å^2^; C) the same as in A) where all structure factors in the range 2-5 Å-resolution were artificially assigned to be equal to zero; D) a set of structure factors of the resolution in the range 2-4 Å taken with the Gaussian weights increasing for higher-resolution structure factors and calculated with point-atom scattering factors, lower resolution structure factors were assigned to be equal to zero.

In our approach (Urzhumtsev and Lunin, 2022a), to model the distortions in the experimental map, we attribute two individual heterogeneity parameters to every atom. The first one is the standard isotropic atomic displacement parameter *B*_*atom*_. The second parameter is atomic image resolution *D*_*atom*_ which reflects the local resolution of the experimental map around the center of the atom. While the first parameter describes a Gaussian-shape attenuation of the magnitudes of Fourier coefficient with resolution, the second parameter stands for other kinds of their absence or reduction, in particular a sharp cut-off in crystallography. We call these parameters the ‘point’ atomic heterogeneity parameters since they are attributes of an isolated atom, and do not consider its environment. The proposed analytical formulas for calculation of model density values and their derivatives with respect to resolution *D*_*atom*_ allow us to include the parameters {*D*_*atom,n*_, *n* = 1,2, … *N*^*atoms*^} in gradient-driven real-space crystallographic refinement (Urzhumtsev and Lunin, 2022b) like ADP refinement.

We may also estimate the local heterogeneity of a given map estimating heterogeneity parameters in different small regions, referred to as vicinities below. To do so, for a given reference point **r**_*ref*_ in space we define the local heterogeneity parameters values *B*_*vic*_, *D*_*vic*_ obtained under the assumption that all atoms neighboring to **r**_*ref*_ have the same *B*_*atom*_ and *D*_*atom*_ values equal to *B*_*vic*_, *D*_*vic*_. The values *B*_*vic*_, *D*_*vic*_ are determined by minimizing the discrepancy between the calculated and observed maps in a vicinity of the reference point. While any space point can be chosen as a reference one, below we consider only the centers of atoms as the reference points. This allows us to consider the local values *B*_*vic,n*_, *D*_*vic,n*_ found in a vicinity of the atom *n* as preliminary estimates of the point heterogeneity parameters *B*_*atom,n*_, *D*_*atom,n*_ of this atom.

A particular difficulty to estimate the local ADP value *B*_*vic*_ and the local resolution *D*_*vic*_ in a vicinity of a given atom is that both factors blur the density peaks somehow similarly. Thus, a decrease in the height and an increase in the width of the central peak in the atomic image can be described both by increasing *B*_*vic*_ or *D*_*vic*_ values (see fragments B and C in Fig. 1). The major difference between the two effects is that limiting the resolution leads to Fourier series termination ripples (James, 1948). The tests below demonstrate that the proposed approach allows in most cases to separate the influence of these two effects and accurately assign both the local resolution and local uncertainty in the positions of atoms.

Our procedure is not limited to crystallographic electron density maps and can be extended, to maps of other physical fields, in particular to the electrostatic scattering potential maps studied by single-particle cryo-EM.

## 2. Methods

### 2.1. Limited-resolution atomic images

The electron density distribution or an electrostatic scattering potential of an isolated immobile atom in the origin are real-valued spherically symmetric functions. In what follows, we call both ‘atomic density distribution’ and denote by *ρ*^*theor*^(**r**). Their integral Fourier transform *f*(**s**), referred to as the atomic scattering factors, are also real-valued and spherically symmetric. We denote the radial parts of these spherically symmetric functions by an overline, 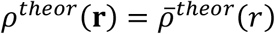 and 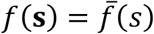, where *s* = |**s**| and *r* = |**r**|. These radial parts are related to each other by the one-dimensional sine Fourier transforms

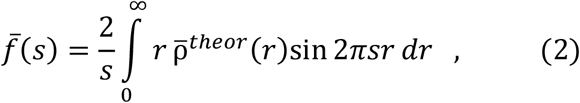

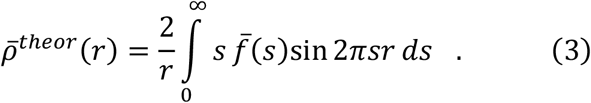

The X-ray and electron scattering factors of all types of atoms and corresponding density distributions are known, as well as their multi-Gaussian approximations (*e.g*., Doyle and Turner, 1968; Peng, 1999)

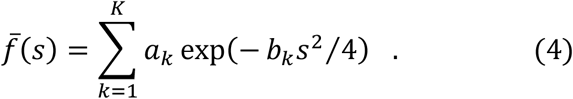

Depending on the required accuracy, the number of terms *K* in these approximations may change (Grosse-Kunstleve *et al*., 2004) and usually varies from two to five.

We define the *D*-resolution image of the atom by integral (3) calculated within the limits 0 ≤ *s* ≤ *D*^−1^, like (1)

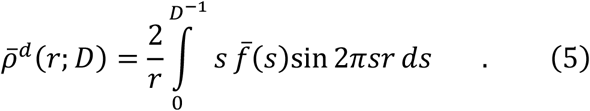

### 2.2. Shell-approximation to the atomic image

The new concept of the shell approximation to atomic image suggested earlier by the authors (Urzhumtsev and Lunin, 2022a, 2022b; Urzhumtsev et al. 2022; Urzhumtseva et al. 2023) contains three key issues. First, it proposes to interpret heterogeneous maps in structural biology by extended atomic models, traditional parameters of which, such as coordinates and displacement parameter, are completed by a point atomic resolution. Second, the atomic images composing such heterogeneous maps are expressed by an analytic function of all these parameters which makes it easy to adjust their values in order to obtain the best fit of the model map to the experimental one. Third, the values of this extended list of parameters can be searched in real space with no direct use of diffraction data. It was shown that for an atom centered at the point **r**_*atom*_, with scattering factor (4), and with the isotropic atomic displacement parameter *B*, its image calculated at the resolution *D* may be represented in any point **r** of space as

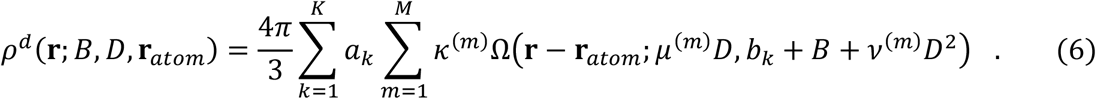

All terms of this sum are expressed through the same spherically symmetric shell-function of a vector **r**

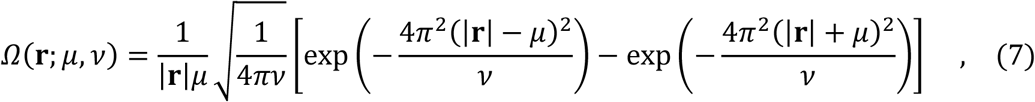

which describes a density distributed uniformly in an ultra-thin spherical shell of the radius *μ* and blurred with the Gaussian function which ‘width’ is characterized by atomic displacement parameter *v*. The resolution value *D* can vary from one atom to another and is associated with the atom, similar to *B*. All parameters of expression (6), except the atomic coordinates **r**_*atom*_ and values *B* and *D*, are known constants. The parameters *a*_*k*_, *b*_*k*_ define the atomic scattering factor (4) and their values may be found, *e*.*g*., in (Peng, 1999) or in (Brown *et al*., 2006). The values *μ*^(*m*)^, *v*^(*m*)^, κ^(*m*)^ are constants of the decomposition of the three-dimensional interference function into a sum of Ω-functions (Urzhumtsev and Lunin, 2022a). An essential feature of decomposition (6-7) is that *D* is now considered a continuous parameter and not a fixed cutoff.

Formulas (6-7) make it possible to calculate a model map adapted to the errors of the experimental map, provided that the parameters of the model atoms are known. We call *B, D* as point heterogeneity parameters of the atom to emphasize that they are related to an isolated atom and do not consider its environment.

### 2.3. Local map heterogeneity parameters

The formulas (6-7) describe morphing the image of an isolated atom when the heterogeneity parameters change. In a real structure, each map value is a sum of the contributions of several neighboring atoms, which can have different values of point heterogeneity parameters. For a point **r**_*ref*_ in space, we define its local map heterogeneity parameters as the values which minimize the discrepancy between the map calculated from the model and the experimental one hypothesizing that the point heterogeneity values are the same for all neighboring atoms. In this work, we choose **r**_*ref*_ coinciding with atomic centers and compare the calculated local map parameters with the point parameters of the heterogeneity of the respective atoms. We formalize this concept as follows.

Let suppose that an experimental map is defined by its values 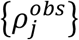 in a grid and the atomic model is given by the coordinates of atoms {**r**_*atom,n*_}. For a point **r**_*ref*_ we define its vicinity *V*_*ref*_ (**r**_*ref*_, *R*_*vic*_) as a sphere of a given radius *R*_*vic*_ centered in **r**_*ref*_. Theoretically, all atoms should be taken to calculate the *V*_*ref*_(**r**_*ref*_, *R*_*vic*_) map values by (6-7). In practice, functions (6) are assigned to be zero beyond a certain distance *R*_*cut*_ from the atomic center, so that only the atoms distanced from **r**_*ref*_ by less than *R*_*cut*_ + *R*_*vic*_ contribute to the density values. At this stage, we suppose that all the atoms contributing to *V*_*ref*_(**r**_*ref*_, *R*_*vic*_) have the same values *B*_*trial*_, *D*_*trial*_ of the point heterogeneity parameters, and calculate by (6-7) the values 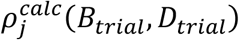 of the model map for the grid nodes in *V*_*ref*_ (**r**_*ref*_, *R*_*vic*_). Then we calculate the discrepancy of the model map values from the experimental ones

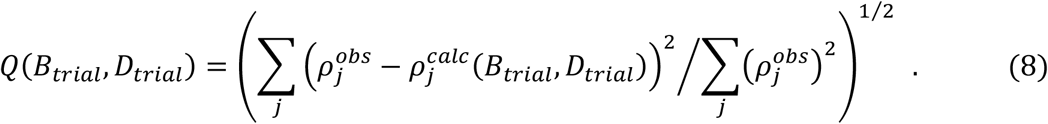

Here the sum is calculated over all grid points which belong to the vicinity *V* (**r**_*ref*_; *R*_*vic*_). We define the local map heterogeneity parameters *B*_*vic*_(**r**_*ref*_), *D*_*vic*_(**r**_*ref*_) for the reference point **r**_*ref*_ as *B*_*trial*_, *D*_*trial*_ values that minimize this discrepancy. If the reference point coincides with the center of *n*-th atom as we do it in this work, i.e., when **r**_*ref*_ = **r**_*atom,n*_, we simplify the notations of the heterogeneity values *B*_*vic*_(**r**_*ref*_), *D*_*vic*_ (**r**_*ref*_) as *B*_*vic,n*_, *D*_*vic,n*_.

### 2.4. Scaling the calculated map

The values to be compared with the least-squares targets such as (8) are often given in different scales. When a linear scaling is acceptable, its parameters κ, *ρ*^0^, unknown *a priori*, can be defined for the considered vicinity together with *B*_*vic*_, *D*_*vic*_ by minimization of the discrepancy

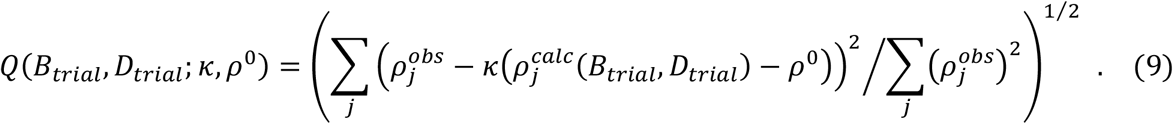

This increases the number of parameters describing the heterogeneity of the experimental map by two individual local scaling coefficients per each vicinity. Eventually, this may overfit the data and lead to instability of the found parameters (Sec. 3.2.2). As a compromise, a two-step procedure has been suggested. In the first run, the map-scaling parameters were determined independently for the vicinity of every reference point. In the second run, the values of parameters κ, *ρ*^0^ were fixed being taken equal to their average values found from the first run.

Often, the input experimental map *ρ*^*obs*^(**r**) is given in the σ-scale, with the zero mean value and a unit root-mean-squares deviation. In this case *ρ*^0^ can be estimated as the total scattering content of the cell normalized by the cell volume. With such complementary information, the first run may be performed with the fixed value of *ρ*^0^ and the only free scale parameter κ.

### 2.5. Test’s protocols

Below, we discuss the results of the tests conducted to find out to what extent the found values of local map heterogeneity parameters reproduce the point values of these parameters for individual atoms. The input data for each test were:

- a density map for the heterogeneity analysis defined in a grid;
- coordinates of the available atomic model corresponding to the map;
- a set of reference points in the vicinity of which the parameters of local map heterogeneity were searched for; in the tests below, they always corresponded to atomic positions;
- control values of the atomic heterogeneity parameters for the reference set of atomic positions; these values were obtained by previous experimental projects or were assigned artificially for the goals of the tests; they were never used to determine the parameters of local map heterogeneity but only to analyze the results.

Three modes of scaling in (9) were used in different runs: fixed κ and *ρ*^0^ values; free κ and *ρ*^0^; free κ and fixed *ρ*^0^.

The method, used in this work, defines the values of the parameters *B*_*vic*_, *D*_*vic*_ by a systematic search on a regular grid by parameters *B*_*trial*_, *D*_*trial*_ minimizing this discrepancy (9). If necessary, the optimal values of one or two scaling coefficients were determined analytically, since the square of the function (9) is a quadratic function of the parameters κ and κ*ρ*^0^.

Below, we present the test results for five different density maps. The tests were done with different scaling protocols and the results are shown for different subsets of atoms (*e*.*g*., the main chain or the side chains atoms). To simplify the navigation, title of each diagram mentions the analyzed map, the main details of the protocol (if necessary), and a set of atoms for which the results are presented.

The program *Pymol* (Schrodinger and DeLano, 2000) was used for illustrations.

### 2.6. Test objects

#### 2.6.1. Crystal structure of initiation factor 2 (IF2)

The suggested approach was evaluated using the structure of the core of the Translation Initiation factor 2 protein from *Thermus thermophilus*, referred below to as IF2. The protein has been crystallized in space group P2_1_2_1_2_1_ with the unit cell dimensions 45.42 × 61.46 × 162.40 Å. The crystal structure (PDB entry 4b3x) has been determined by MAD and refined with the program *phenix.refine* (Afonine *et al*., 2012) at the resolution of 1.95 Å (Simonetti *et al*., 2013) with a near complete diffraction data set. The atomic model extracted from the PDB (Burley *et al*., 2021) contained 2908 atoms of the protein, 220 atoms of water molecules and a glycerol molecule. The protein molecule consists of four domains with a degree of order variable from clearly visible alternative conformations for some residues to highly disordered loops and the C-terminal region. This variability is reflected by the values of its atomic displacement parameters (Fig. 2). These values extracted from the PDB atomic model (*B*_*atom*_ below) have never been used for the heterogeneity analysis but only to validate the results of the tests. The suggested method was applied first to the maps calculated with simulated IF2 data and then to two maps calculated using experimental structure factor magnitudes as explained below.

**Fig. 2.**
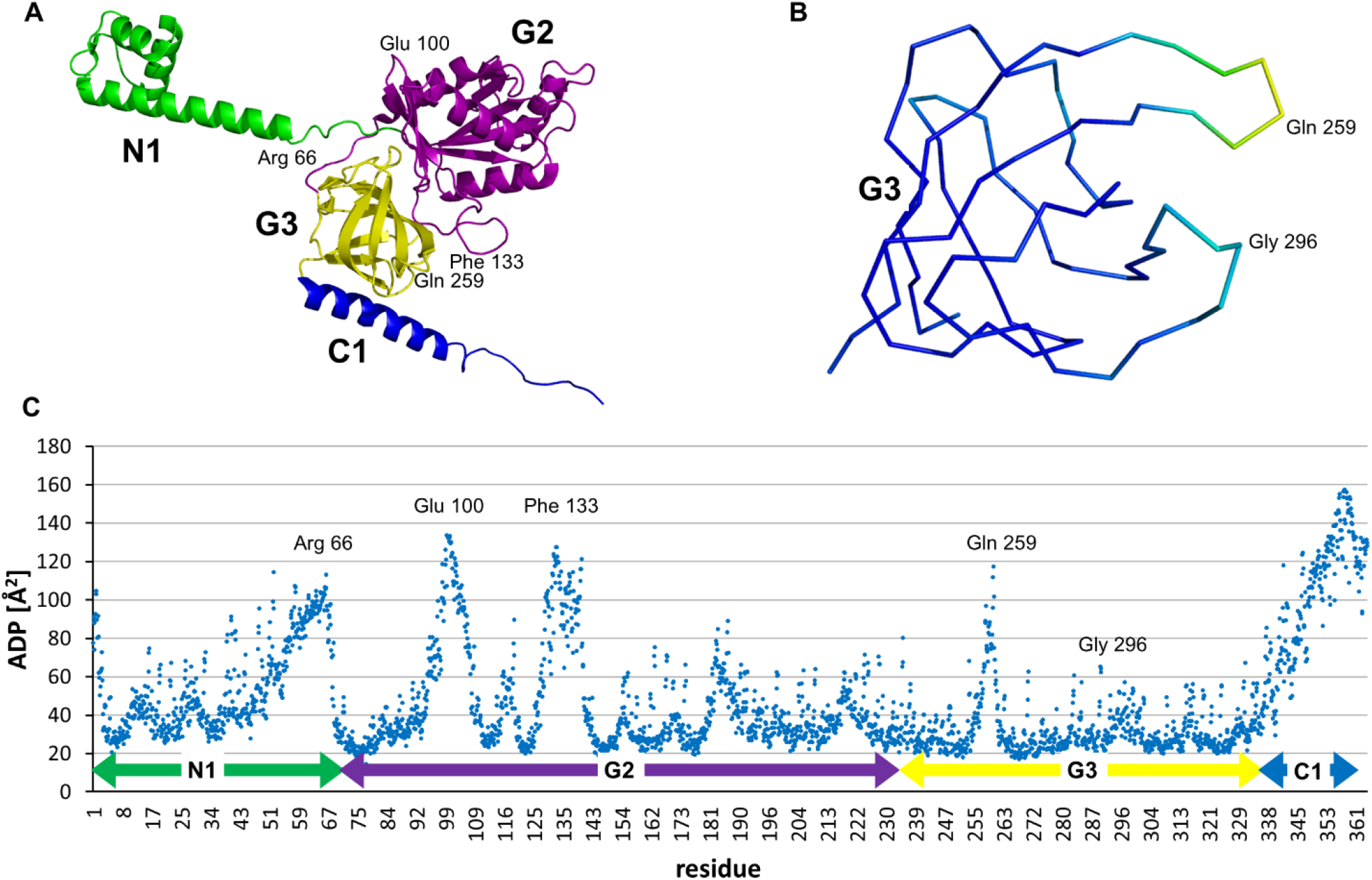
A. Atomic model of Initiation Factor 2 and its domains. B. Domain G3 backbone colored according to the *B*-factor. C. Distribution of the *B*-factor values in the IF2 model.

#### 2.6.2. Simulated single particle domain G3 in a virtual unit cell (G3_P1_)

To avoid an influence of various experimental, computational, and human errors, we started from the tests with simulated maps. Each such map was calculated as a sum of atomic images, each with its own assigned values *B*_*atom*_, *D*_*atom*_ of atomic displacement and atomic image resolution parameters.

In the first test we considered the domain G3 (83 residues; 682 atoms including 345 main chain atoms N, C_α_, C, O). This domain contains both regular parts and highly disordered loop 357-361 poorly visible in the maps (Fig. 2). Except for the disordered loop, the ADP values *B*_*atom,n*_ for the main chain atoms are in the range 20-25 Å^2^, reaching significantly higher values for side chains (Fig. 2C).

The model was placed in the center of a virtual orthogonal unit cell of the size 54.0 × 54.0 × 48.0 Å in the space group P1 (G3_P1_ model). It had no contacts with its translationally-related copies simulating a virtual crystal considered in single-particle studies. The values {*B*_*atom,n*_} were extracted from PDB and all values {*D*_*atom,n*_} were assigned equal to 2.0 Å. These values were considered as the control values when analyzing the test results. The map G3_P1_was calculated in a grid with the step 0.5 Å as the sum of 2.0 Å-resolution images of the model atoms taken in the absolute scale (*e*Å^−3^).

#### 2.6.3. Simulated single particle heterogeneous resolution map (IF2_VR_)

For the second series of tests, a single copy of the whole IF2 model was placed at the center of a virtual orthogonal unit cell of the size 80.0 × 120.0 × 100.0 Å in P1 space group. As in the previous test, the molecule had no contacts with its translationally-related copies and the {*B*_*atom,n*_} values were taken from PDB. However, this time the value *D*_*atom,n*_ assigned to different atoms varied from 2 Å for the atoms in the central part of the molecule, inside the sphere of the radius 10 Å, to *D*_*atom,n*_ = 5 Å for the atoms outside the sphere of the radius 40 Å, increasing linearly with the distance to the center in between. The corresponding map was calculated as the sum of atomic images of the respective resolution in the grid with the step 0.5Å. The same map was taken in different scale in two tests: in the absolute scale or in the σ-scale. This latter simulated the practical situation when the absolute scale of the map is unknown.

#### 2.6.4. IF2 crystallographic maps (IF2_obs_ and IF2_σA_)

Two experimental crystallographic maps of 2 Å resolution were calculated with coefficients downloaded from the PDB (4b3x). The first map, IF2_obs_, was calculated with the experimentally observed structure factors magnitudes *F*^*obs*^ and the phases calculated from the refined atomic model (Simonetti et al., 2013). Differently from the previous maps calculated with simulated data, this map had extra imperfections due to magnitude and phase errors. The goal of the test was to estimate to what extent all map distortions can be modeled by only two types of local map heterogeneity parameters {*B*_*vic,n*_} and {*D*_*vic,n*_}, and how well these parameters match the parameters {*B*_*atom,n*_} extracted from PDB and the formal resolution parameters *D*_*atom,n*_ = 2Å.

The second map, IF2_σA_, was the *σ*_*A*_-weighted map (Read, 1986) in which the magnitudes are weighted combinations of the experimental and model ones reflecting estimated accuracy of the suggested phases. Such weights are expected to improve the quality of the map, but may be a source of additional map distortions modifying the magnitudes. The goal of the test with such map was to check the possibility to describe additional distortions by the same two local map heterogeneity parameters.

The both maps were calculated at the formal resolution 2.0Å in the grid with the step 0.5Å along each coordinate axis.

#### 2.6.5. Cryo-EM structure of human ribosome (HR)

Finally, the proposed approach was tested with an experimental cryo-EM map of the human ribosome, HR (Frechin *et al*., 2023), kindly provided by the authors. In that original study, the initially found map of an estimated resolution of 2.28 Å was refined and sharpened using the *B* factor filtering of -41.11 Å^2^. This increased the resolution up to approximately 2.18 Å. The local resolution was estimated by the Fourier shell correlation (FSC) with the 0.143 cut-off done by Relion (Zivanov *et al*., 2018). Map interpretation was done by docking the atomic model of the 80S human ribosome (PDB ID 6QZP) (Wang *et al*., 2021) into the final maps followed by an automated rigid body fit with Chimera (Pettersen *et al*., 2004), manual rebuilding with Coot (Emsley *et al*., 2010) and real space refinement with Phenix (Afonine *et al*., 2018). This provided refined atomic coordinates and individual {*B*_*atom,n*_} and {*D*_*atom,n*_} values. Fig.3 shows the fragment of the available atomic model, Lys 43 – Tyr 73 of the chain Lh, superposed with the experimental map. This map shows unambiguously the main chain atoms and the most of side chains, but not all.

**Fig. 3.**
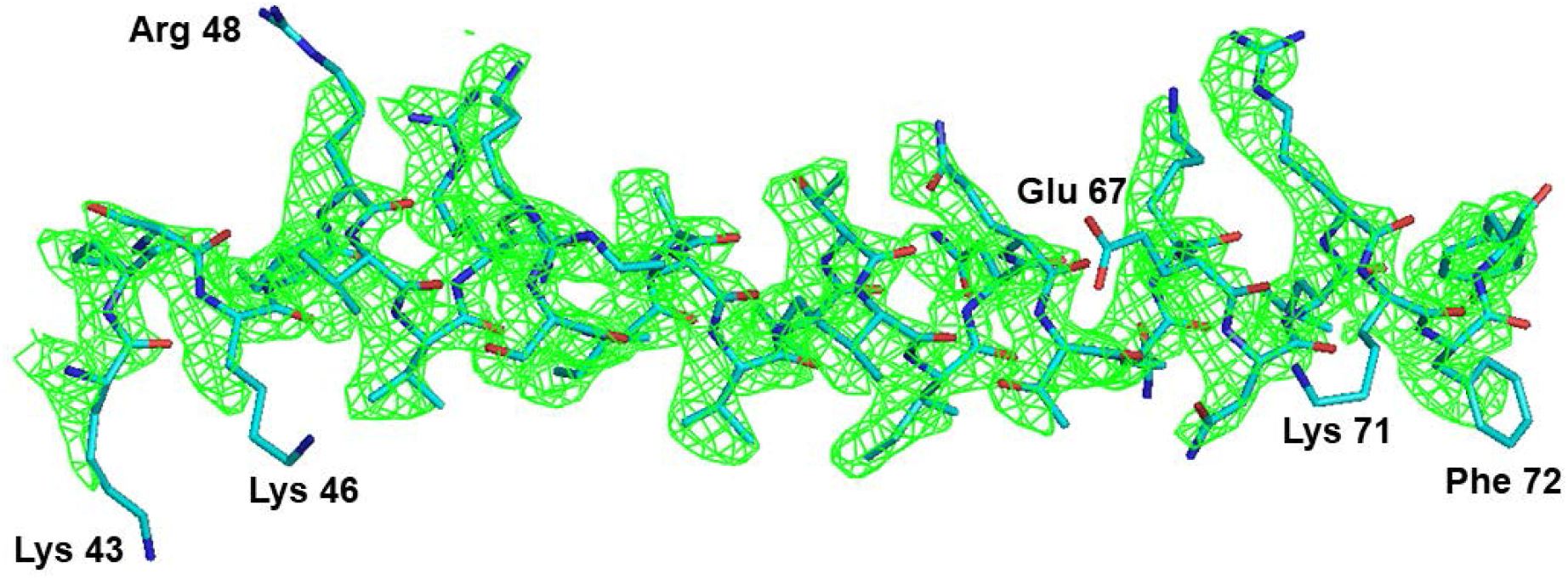
The fragment Lh Lys 43 – Tyr 73 of the human ribosome superposed with the experimental cryo-EM map.

## 3. Results

### 3.1. G3_P1_ homogeneous resolution map

The goals of this test were to check the ability of the method to separate the influence of the displacement and resolution parameters on the molecular image in the map, to estimate if the calculated map heterogeneity values *B*_*vic*_, *D*_*vic*_ can be used as an approximation to the point atomic heterogeneity values *B*_*atom*_, *D*_*atom*_ and to study an eventual variation of the estimated local resolution parameters *D*_*vic*_ for a map of a homogeneous resolution. The map was calculated in the absolute scale (*e*Å^-3^) as the exact image of the G3 domain placed in a virtual unit cell in space group P1 (see Section 2.6.2), using {*B*_*atom,n*_} values extracted from PDB file and all {*D*_*atom,n*_} values assigned as 2 Å. The set of atomic coordinates extracted from PDB was used both as coordinates of the model atoms **r**_*atom,n*_ and as the reference points {**r**_*ref,n*_}. The trial values of both *B*_*vic*_ and *D*_*vic*_ varied during the search with the steps 10.0 Å^2^ and 0.5 Å, respectively. The vicinity radius and density cutoff radius were taken as *R*_*vic*_ = 2.1 Å and *R*_*cut*_ = 3*D*_*trial*_ Å, depending on the trial resolution value (Urzhumtsev et al., 2022).

Fig. 4 shows the deviation of the found *B*_*vic,n*_ values, obtained in this test, from the control atomic *B*_*atom,n*_ values. For most of atoms the difference between the *B*_*vic,n*_ and *B*_*atom,n*_ was within the single search step. The mean absolute value of the difference was 3.27 Å^2^ for the whole domain.

**Fig. 4.**
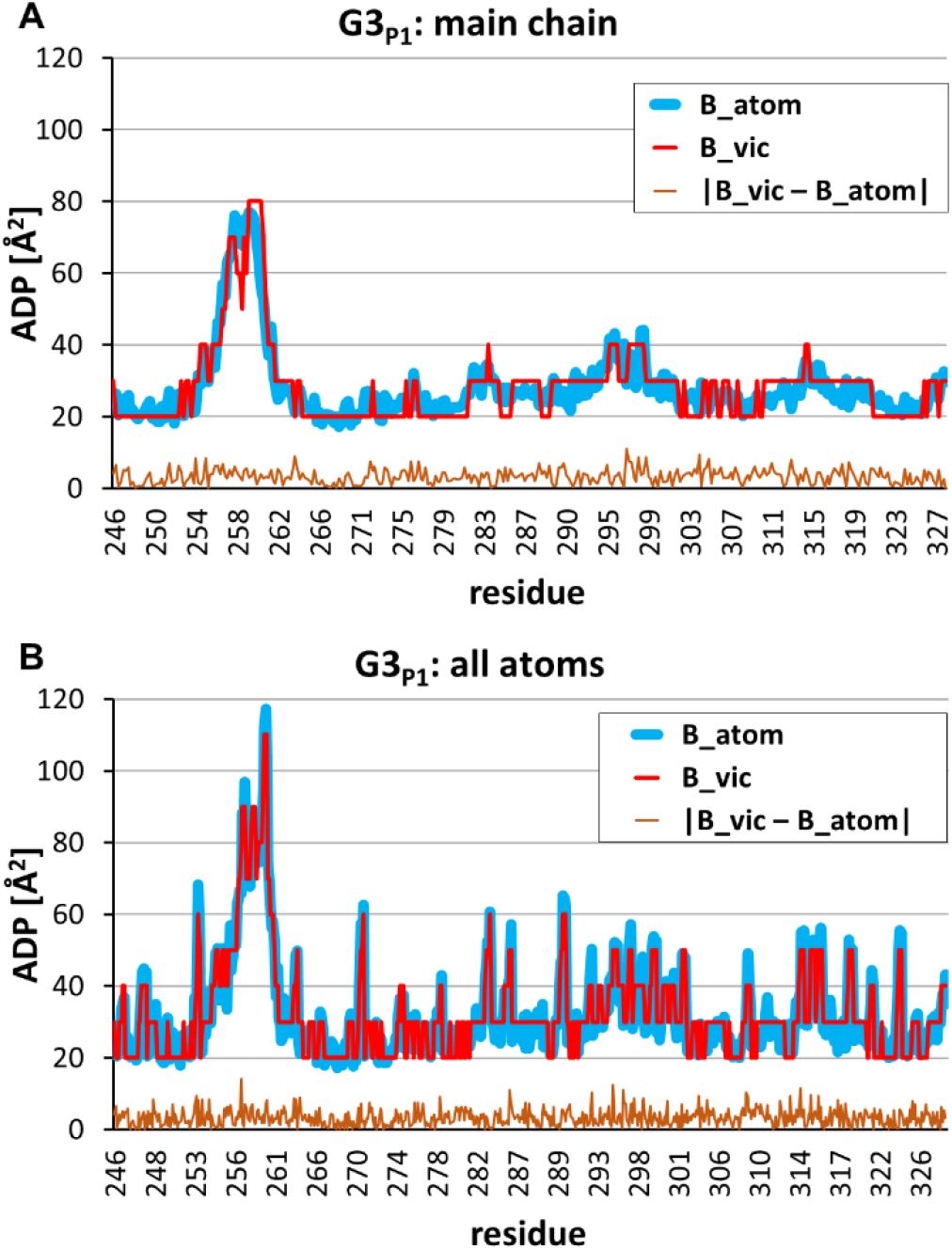
Results of the heterogeneity analysis of the G3_P1_ map of the 2 Å-resolution. The values *B*_*vic*_ were searched with the step of 10 Å^2^. The *B*_*vic*_ values, the control *B*_*atom*_ values, and their absolute difference are shown for the main chain atoms (A) and for all atoms (B).

For a vast majority of atoms, the value of *D*_*vic*_ was found to be equal to *D*_*atom,n*_ = 2.0 Å used to calculate the map. The only exceptions were observed for a few *D*_*vic*_ values equal to 2.5 or 1.5 Å, just neighboring to 2.0 Å. This occurred for 25 from the total number of 682 atoms, including 5 main chain atoms from 345, and corresponded to the mean absolute deviation of *D*_*vic,n*_ from *D*_*atom,n*_ equal 0.02Å for all atoms (no plot shown).

This test established two important facts. First, the suggested method makes it possible to separate the influence of ADP and AIR on the distortion of the atomic image in the map. Second, when being applied to the map of a homogeneous resolution, the given freedom to change the value of the AIR parameter is not taken and, as a rule, the correct value is chosen.

### 3.2. Heterogeneity analysis of IF2_VR_ maps of a variable resolution

#### 3.2.1. Analysis of the map in the absolute scale

The goal of the next test was to check the ability of the procedure to estimate individual atomic image resolution values for maps of variable resolution. The analyzed map was calculated in the absolute scale (*e*Å^-3^) as the exact image of IF2_VR_ atomic model with simulated variable atomic image resolution values. The model was placed in a virtual unit cell, the {*B*_*atom,n*_} values were extracted from PDB file, and the {*D*_*atom,n*_} values were assigned as described in Sec. 2.6.3. The set of atomic coordinates extracted from PDB was used both as coordinates of the model atoms {**r**_*atom,n*_} and the reference points {**r**_*ref,n*_}. The trial values of both *B*_*vic*_ and *D*_*vic*_ varied during the search with the steps 5.0 Å^2^ and 0.2 Å, respectively. The vicinity radius and density cutoff radius were chosen as *R*_*vic*_ = 2.1 Å and *R*_*cut*_ = 3*D*_*trial*_ Å.

Fig. 5 shows the results of the test. For the most of atoms, the difference between the found *D*_*vic*_ and the control *D*_*atom*_ values was within a single search step. The mean absolute difference between these values was equal to 0.10 Å for the main chain atoms and 0.15 Å for the side chain atoms. The mean absolute difference between *B*_*vic*_ and *B*_*atom*_ was 5.40 Å^2^ for the main chain atoms and 6.88 Å^2^ for the atoms in side chains. However, for some atoms, especially for those with large *B*_*atom*_ values or in the low-resolution parts of the map (Fig. 5E), this deviation was larger. It is worthy of noting that in poor defined regions of the map, the found values *D*_*vic,n*_ were sometimes larger and sometimes smaller than the respective *D*_*atom,n*_ values. When the found *D*_*vic,n*_ > *D*_*atom,n*_, the corresponding value *B*_*vic,n*_ was smaller than *B*_*atom,n*_ (upper left corner in Fig. 5F) to conserve the degree of blurring of the peak in the map. Inversely, for atoms with *D*_*vic,n*_ < *D*_*atom,n*_ the corresponding value *B*_*vic,n*_ was larger than *B*_*atom,n*_ (bottom right corner in Fig. 5F).

**Fig. 5.**
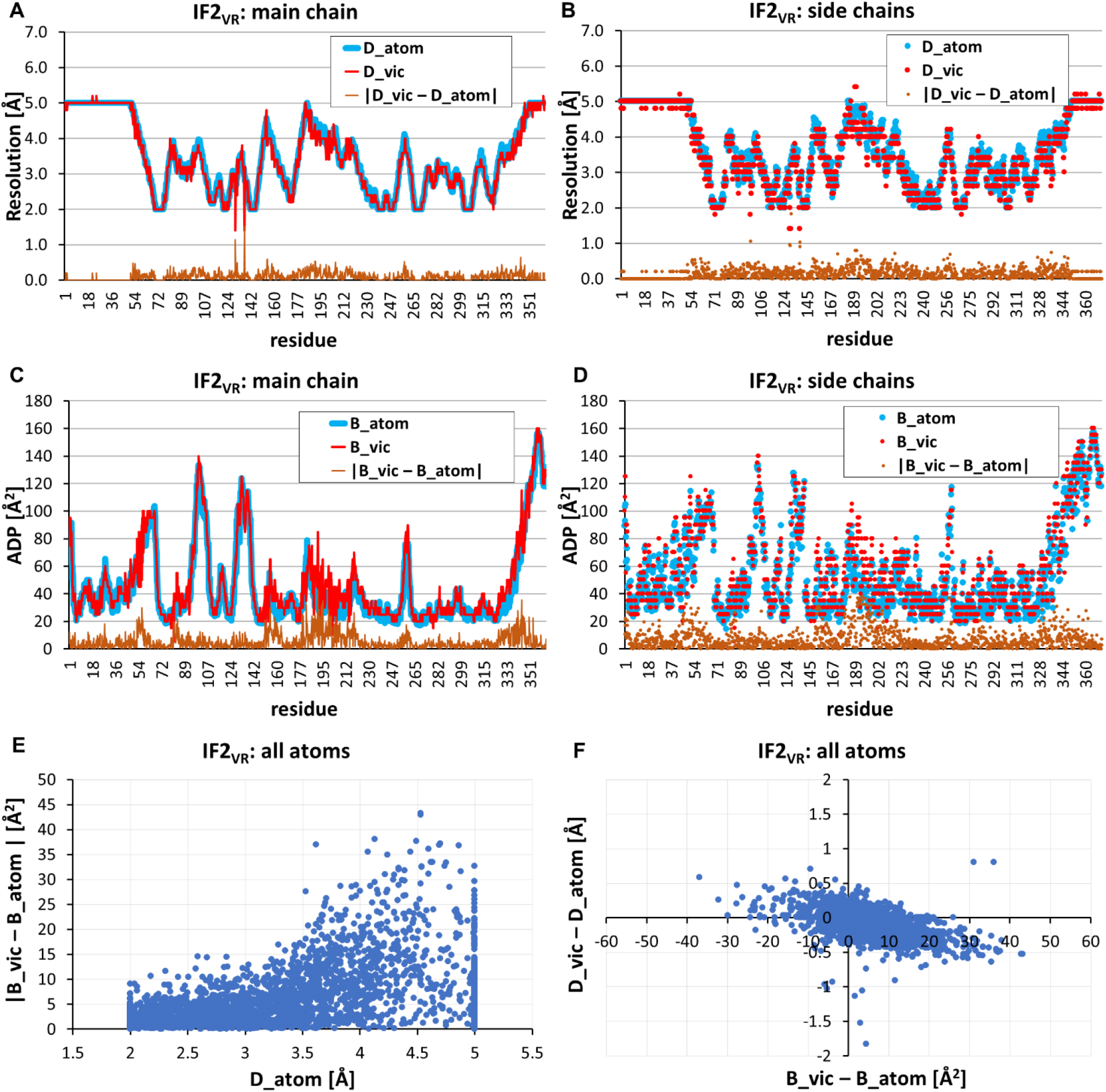
Results of the heterogeneity analysis of the IF2_VR_ map of a variable resolution. The values *B*_*vic*_ were searched with the step of 5Å^2^, and *D*_*vic*_ the with the step of 0.2 Å. A, B) The *D*_*vic*_ values, the control *D*_*atom*_ values, and their absolute difference for the main chain and side chains atoms. C, D) The *B*_*vic*_ values, the control *B*_*atom*_ values, and their absolute difference for the main chain and side chain atoms. E) Joint distribution of the absolute deviation of *B*_*vic*_ from *B*_*atom*_ and *D*_*atom*_ value plotted for all atoms; each point in the diagram corresponds to one atom. F) Joint distribution of the deviation of *D*_*vic*_ from *D*_*atom*_ and the deviation *B*_*vic*_ from *B*_*atom*_ plotted for all atoms; each point in the diagram corresponds to one atom.

#### 3.2.2. Heterogeneity analysis of a normalized map

In the test above, both the analyzed map *ρ*^*obs*^(**r**) (*e*Å^-3^) and the atomic images for the trial maps *ρ*^*calc*^(**r**) were calculated in the same, absolute scale (*e*Å^-3^) simplifying the search. In practice, maps are usually known in an arbitrary scale and often reduced to the σ-scale. As a result, to perform the search for local heterogeneity parameters by minimization of (9), the trial map, calculated initially in the absolute scale, should be linearly scaled in (9), to match the scale of the observed map. The two scaling coefficients κ (dimensionless) and *ρ*^0^ (*e*Å^−3^) may be considered as additional parameters of the search by minimization of (9), *i.e*., four parameters 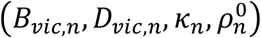 are now defined for the vicinity of every reference point (see Methods).

To simulate such situation, a map was calculated in the absolute scale (*e*Å^-3^) from the IF2_VR_ atomic model placed in a virtual unit cell, using {*B*_*atom,n*_} values extracted from PDB file and with {*D*_*atom,n*_} values assigned as described in Section 2.6.3. Then the map was recalculated in the σ-scale with the coefficients 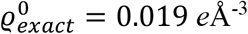 and κ_*exact*_ = 9.*7*41. The set of atomic coordinates extracted from PDB was used both as coordinates of the model atoms {**r**_*atom,n*_} and the set of reference points **r**_*ref,n*_. The trial values of both *B*_*vic*_ and *D*_*vic*_ varied during the search with the steps 5.0 Å^2^ and 0.2 Å, respectively. The vicinity radius and density cutoff radius were taken equal to *R*_*vic*_ = 2.1 Å and *R*_*cut*_ = 3*D*_*trial*_ Å. Figs. 6, 7 show the results of test performed with different scaling protocols.

**Fig. 6.**
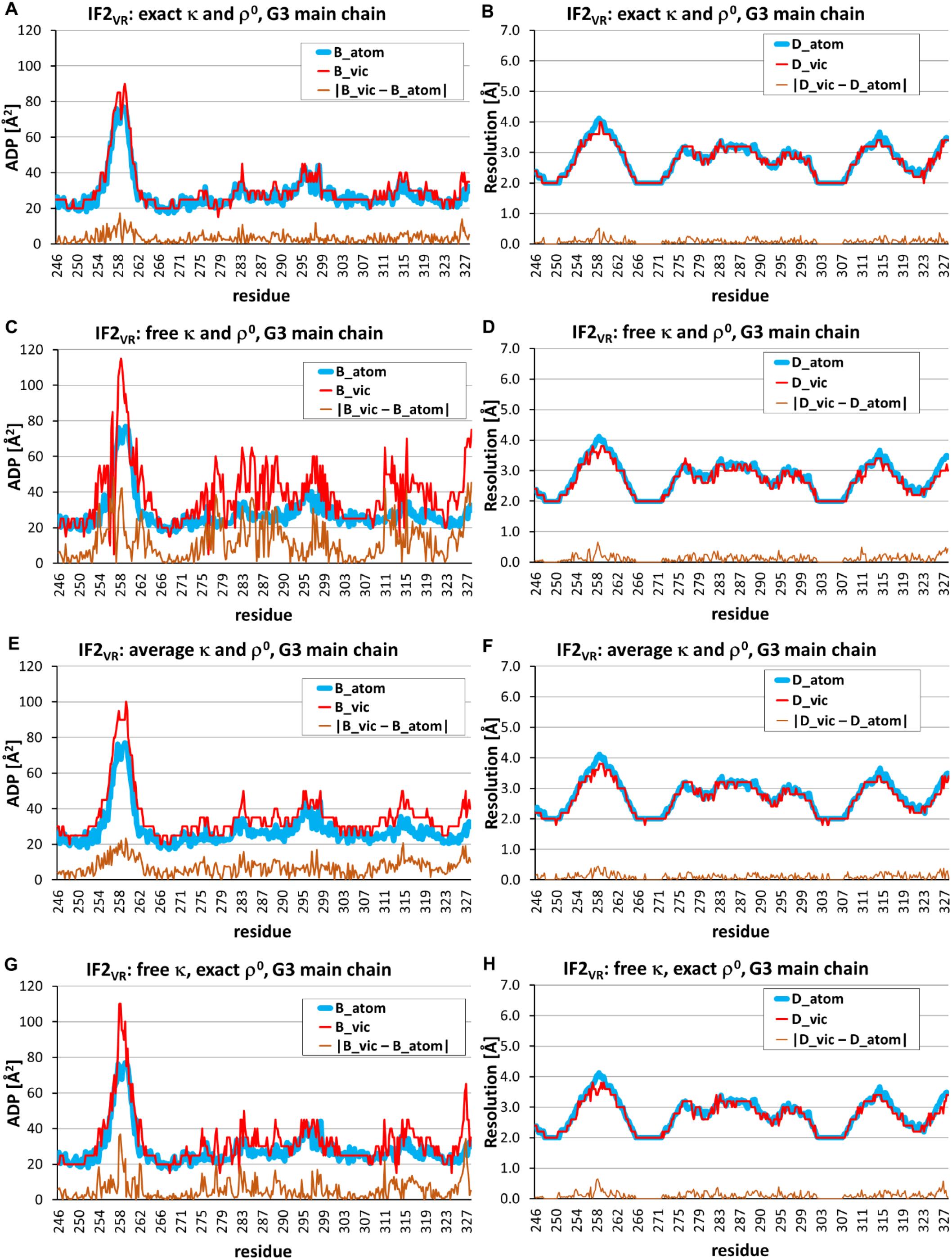
Results of the heterogeneity analysis of the σ-normalized map for the IF2_VR_ map of a variable resolution using different search protocols. The results for the main chain atoms in G3 domain are shown. A, B) The exact scale values κ and *ρ*^0^ applied in a vicinity of each atom; C, D) scale values adapted individually in the vicinity of each atom; E, F) scale values taken in the vicinity of each atom as the average values found from the previous run. G, H) the exact *ρ*^0^ value fixed for the vicinity of each atom while κ value adapted individually in each vicinity. Left column shows model values *B*_*atom*_, found values *B*_*vic*_, and their absolute deviations. Right column shows the same for the *D*_*atom*_ and *D*_*vic*_values.

**Fig. 7.**
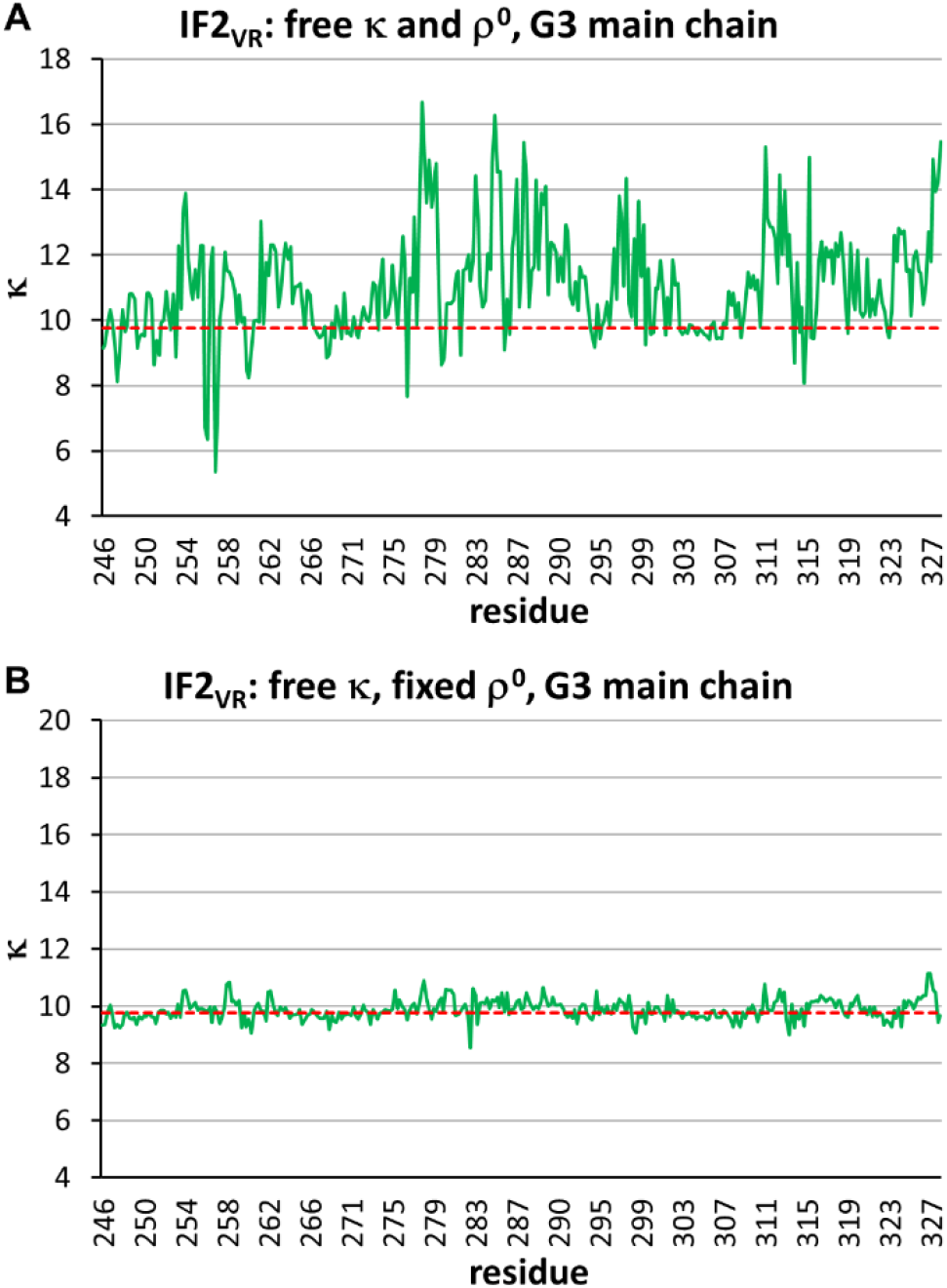
The value of the individual scale factor κ_*n*_ in the vicinity of the main-chain atoms of the G3 domain of IF2_VR_; the values are found from the analysis of the σ-normalized map of a variable resolution. A) Search with variable κ and *ρ*^0^ values; B) search with variable κ and fixed *ρ*^0^ value equal to the exact value. Dashed lines show the exact parameter value known in this test case.

In the first run, the scaled coefficients were fixed to the exact values *k*_*n*_ = *k*_*exact*_, 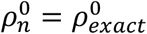.

The results were similar to those from the previous test, that performed in the absolute scale. The mean absolute difference of *B*_*vic*_ and *B*_*atom*_ was 6.13 Å^2^ for all atoms of the model, and the mean absolute difference of *D*_*vic*_ and *D*_*atom*_ was 0.12 Å (Fig. 6A, B). The second test was performed assuming that scale coefficients are unknown, and they were searched for every atom independently, by minimization of (9) (Fig. 6C, D). The results of the second test were worse and illustrated that overparameterization leads to instability in values of both *B*_*vic,n*_ and κ_*n*_ (Fig. 6 C,7A). The mean absolute difference of *B*_*vic*_ and *B*_*atom*_ has increased to 14.68 Å^2^. Surprisingly, the difference between *D* values remained at the same level of 0.13 Å. Then the values κ_*n*_, 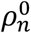 found for all atoms were averaged, resulting in κ 10.3*7*3 and 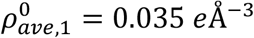. These average values were taken as fixed scaling coefficients in the next run (Fig. 6E, F). Now the absolute difference between the control and determined parameter values was 14.96 Å^2^ for *B* and 0.24 Å for *D*. As an alternative scaling protocol, we fixed the 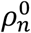 values making them equal to the value calculated analytically from the atomic content of the virtual crystal and equal to the 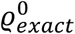 value, used in the map normalization. The run performed with such 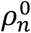 values (Fig. 6G, H), resulted in the absolute difference for map and atomic B and *D* values equal to 9.92 Å^2^ and 0.13 Å, respectively. The κ_*n*_ values varied much less (Fig. 7B) and their average κ_*ave*,2_ = 9.*7*39 was very close to the exact κ_*exact*_ = 9.*7*41 value used in the map normalization. The *B*_*vic*_ and *D*_*vic*_ found with κ_*ave*,2_, 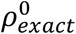 were as accurate as in the test with the exact scaling coefficients (Fig. 6A, B).

To check the role of the search step, the calculation was repeated with fixed values of the scaling coefficients κ_*ave*,2_, 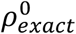 and the steps of the trial values increased to 10.0 Å^2^ for *B*_*vic*_ and to 0.5 Å for *D*_*vic*_. The obtained mean absolute difference between the map and atoms heterogeneity values *B* and *D* for all atoms in G3 domain became 5.27 Å^2^ and 0.16 Å. This shows that such coarser steps 10.0 Å^2^ and 0.5 Å are sufficient to estimate the initial values of local heterogeneity parameters.

### 3.3. Heterogeneity analysis of IF2 crystallographic maps

#### 3.3.1. Unweighted map

Finally, the developed approach was applied to two crystallographic maps of the crystal structure of IF2 for which a near complete set of structure factors and the model refined at resolution of 1.95 Å were available. The first was calculated as the Fourier synthesis (1) with the coefficients *F*^*obs*^(s)*eee*[*iφ*^*calc*^(s)], combining the experimentally obtained magnitudes *F*^*obs*^(s) with the phases *φ*^*calc*^(s) calculated from the model, both available from the PDB. All structure factors of the resolution up to 2.0 Å were included into the synthesis calculation (Section 2.6.4).

First, the analysis was performed with variable map-scaling parameters κ and *ρ*^0^, *i.e*., in each vicinity the calculated density was scaled to the observed one with its individual coefficients. Expectedly from the test calculations, such search has resulted in an unstable behavior of the map-scaling coefficients leading to a relatively large deviation of some found *B*_*vic*_ values from the respective model ADP values *B*_*atom*_ (Fig. 8). Nevertheless, the mean absolute deviation was not so large and constituted 13.97 Å^2^ for all atoms. The mean absolute deviation of *D*_*vic*_ from the actual value *D*_*atom*_ = 2Å, the same for all atoms, was 0.149 Å for the main chain and 0.180 Å for all atoms. The averaging of the found scaling parameters κ and *ρ*^0^ over the main chain atoms of the relatively stable G3 domain resulted in the values 4.08 and 0.347 *e*Å^-3^. The last value is quite close to the estimate *F*_000_/*V*_*cell*_ ≈ 0.363 *e*Å^−3^, calculated from the PDB model content.

**Fig. 8.**
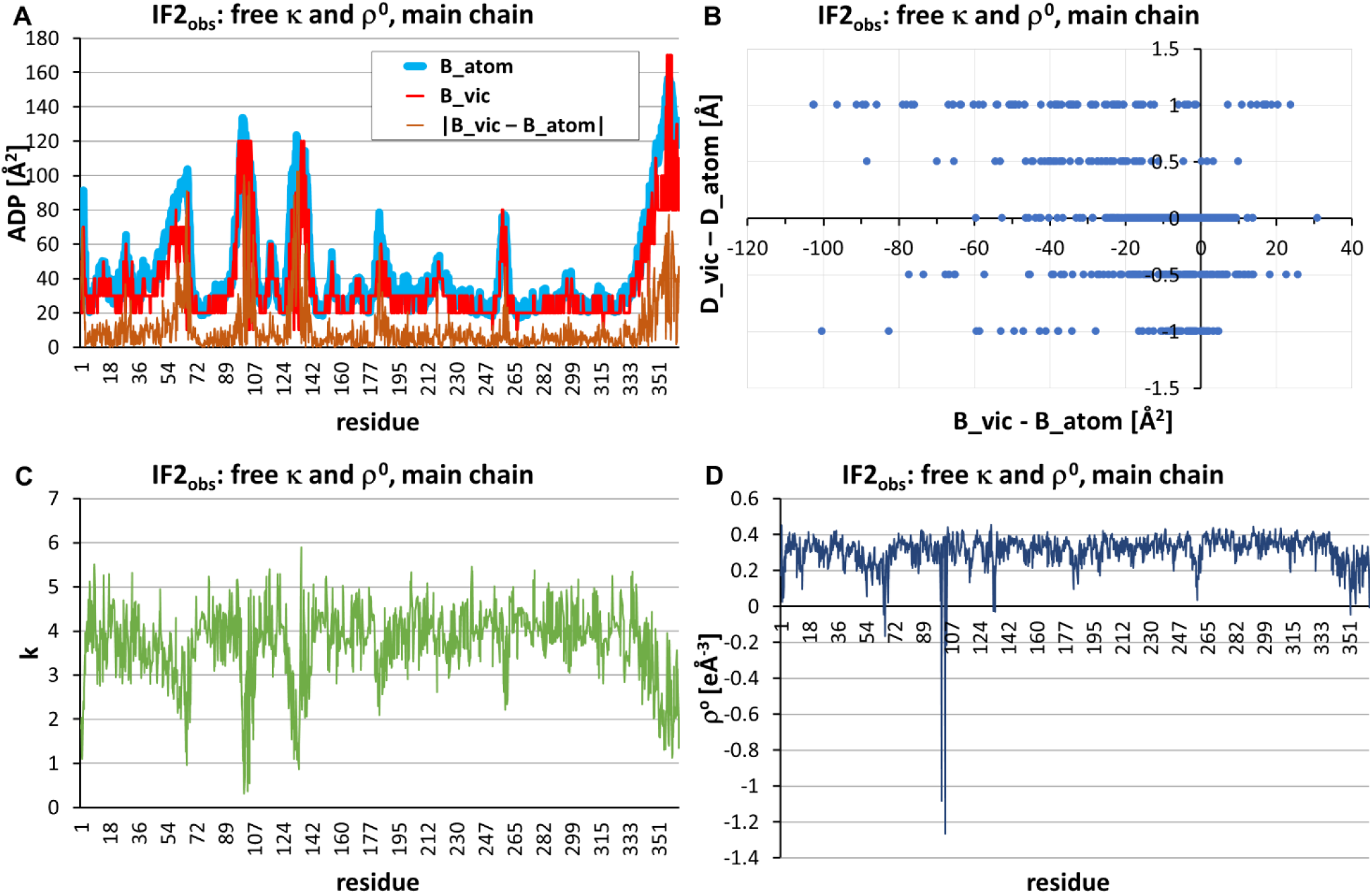
Heterogeneity analysis of the 2 Å-resolution Fourier synthesis for IF2 structure calculated with the coefficients *F*^*obs*^, *φ*^*calc*^. The results are shown for the main chain atoms. The run was performed with the adjustable map-scale parameter values. A) The values *B*_*vic*_, *B*_*atom*_, and their absolute difference. B) The joint distribution of the deviations *D*_*vic*_ − *D*_*atom*_ and *B*_*vic*_ − *B*_*atom*_. C) The found scale factors κ and D) The found scale values *ρ*^0^.

Then, the procedure was repeated taking both the parameters κ and *ρ*^0^ fixed, equal to the mean values found from the first run. Now, the estimated resolution values *D*_*vic*_ coincided with the formal value of 2.0 Å for the most of atoms (Fig. 9 C, D), with the mean absolute deviation equal to 0.29 Å. The deviation of the *B*_*vic*_ values from the control ones was reduced to 7.80 Å^2^. However, while the mean deviation values were small comparatively to the search steps 0.5 Å and 10.0 Å^2^, the errors for some atoms were significant (Fig. 9).

**Fig. 9.**
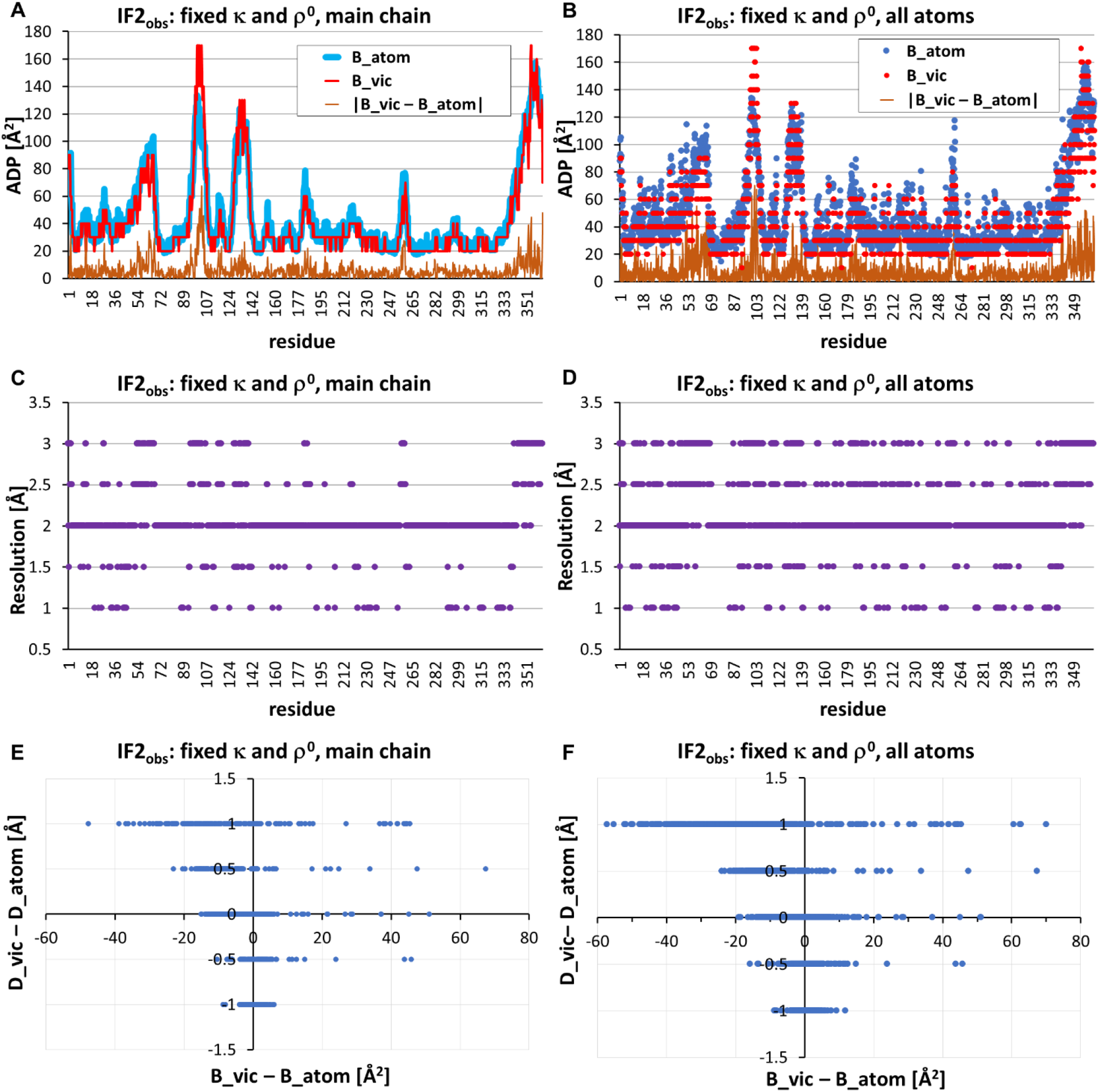
Heterogeneity analysis of the IF2 2 Å-resolution Fourier synthesis, calculated with the coefficients *F*^*obs*^, *φ*^*calc*^. The analysis was done with fixed values of the scaling parameters κ and *ρ*^0^, taken in the vicinity of each atom as the average values found from the previous run. A, B) The values *B*_*vic*_, *B*_*atom*_, and their absolute differences. C, D) The found *D*_*vic*_ values. E, F) The joint distribution of the deviations *D*_*vic*_ − *D*_*atom*_ and *B*_*vic*_ − *B*_*atom*_.

#### 3.3.2. Standard σ_A_-weighted map

The second experimentally-based IF2 crystallographic map was the standard σ_A_-weighted one (Read, 1986) with the coefficients calculated by *phenix.refine* and available in PDB. Similar to the previous calculations, the search for the *B*_*vic*_, *D*_*vic*_ parameters was performed in two iterations, but differently to the analysis of the unweighted map (Sec.3.3.1), now we fixed *ρ*^0^ to the value 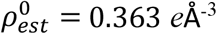 calculated from the PDB model content, while the parameter κ was considered as free and searched for in the first run. The deviations of its found values from the mean value were clearly correlated with the values of the score function *Q* (9) (Fig. 10). By this reason, the average κ-value to be used in the second run was calculated considering only the main-chain atoms with *Q* < 0.2 and has been found as κ_*ave*_ = 4.43. In the second run the values of scale parameters were fixed to κ_*ave*_and *ρ*^0^ values. Fig. 11 shows results of the second run for the atoms of G3 domain. For most of them the values of *B*_*vic*_, *D*_*vic*_ reproduced the values of the corresponding parameters of the atoms as accurately as in the test with the simulated data (Fig. 7G). The number of atoms with the found *D*_*vic*_value different from the formal synthesis resolution 2 Å, was 37 from 345 main chain atoms, and 140 from all 682 atoms. The mean absolute deviation from the formal resolution value 2.0 Å was 0.20 Å for all atoms of the G3 domain, and the mean absolute deviation of *B*-value was 5.14 Å^2^. The plot of the obtained values of the score function (9) (Fig. 11 C) reflects anomalies in the defined *B*_*vic*_, *D*_*vic*_ values which may be caused by insufficient quality of the respective map regions and of the atomic model built there.

**Fig. 10.**
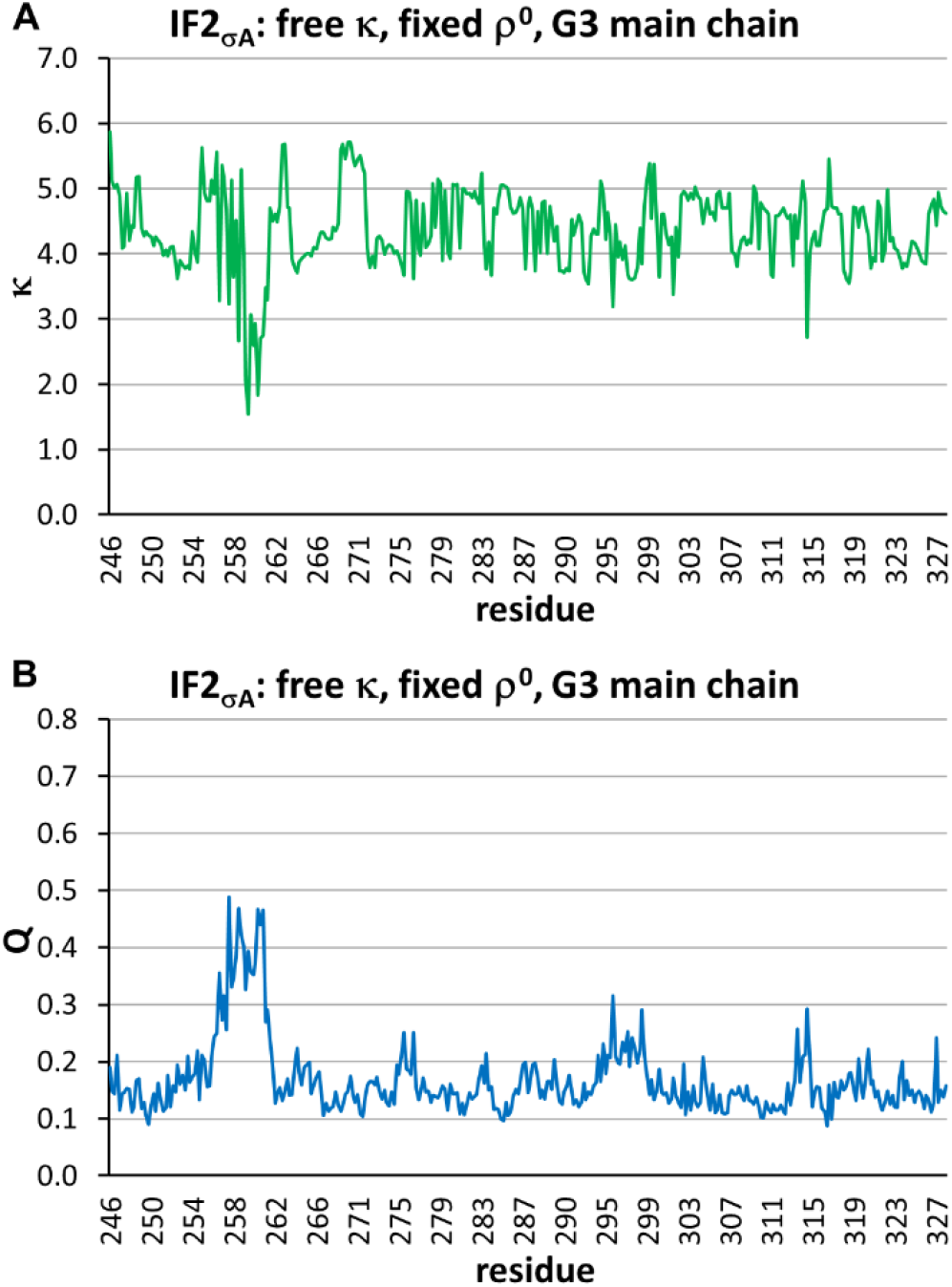
Heterogeneity analysis of σ_A_-weighted map for G3 domain main chain performed with a variable coefficient κ and the fixed *ρ*^0^ value, estimated from the PDB model content. A. Variation of κ_*n*_ along the chain. B. Variation of the score function (9) value along the chain.

**Fig. 11.**
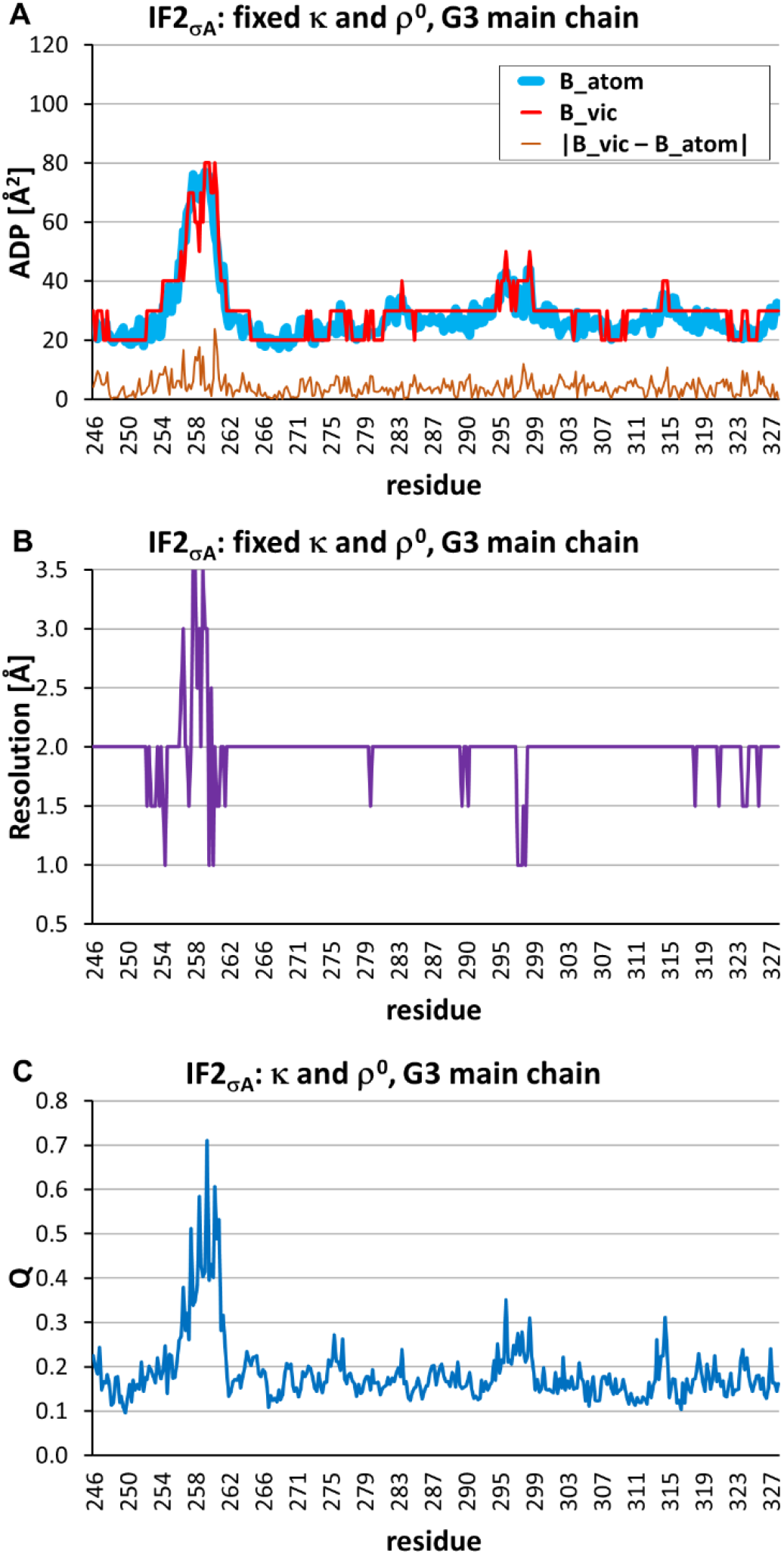
Heterogeneity analysis of σ_A_-weighted map for IF2 with the fixed κ_*ave*_ and 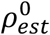 coefficients. The results are shown for the G3 main chain atoms. A. Variation of the *B*_*vic*_, *B*_*atom*_ values along the chain and their absolute difference. B. Variation of the *D*_*vic*_ value along the chain C. Variation of the score function (9) value along the chain.

### 3.4. Heterogeneity analysis of the cryo-EM human ribosome map

The proposed approach could be applied to the maps obtain not necessary by crystallography but by other techniques, in particular by cryo-EM. However, refinement of atomic models against cryo-EM data is not yet so well-established as in crystallography, and its results should be taken with some caution. In particular, cryo-EM maps are often sharpened and postprocessed, so the estimated ADP values may have no clear physical meaning and may vary in the range different from that usual for crystallography. Also, the resolution defined from the FSC analysis determines the reproducibility of the result (precision) rather than the visual quality of the map and may differ from the crystallographic meaning of this term (Penczek, 2010; Afonine *et al*., 2018). For the goals of this project, we used the ribosome experimental data and the respective model as described in Section 2.6.5.

Similar to the previously described cases, the evaluation of the *B*_*vic*_, *D*_*vic*_ parameters for the atomic model was performed in two iterations. In the first run, the scale parameters κ, *ρ*^0^ were considered as variable and defined individually in the vicinity of every atom together with *B*_*vic*_, *D*_*vic*_. Then, the scale parameters defined for main chain atoms were averaged and these average values were fixed and used in the second iteration. Atomic images were calculated using the 4-Gaussian approximation to the atomic scattering factors (Peng, 1999). The radius of the vicinity in these tests was taken equal to 4 Å. The steps in *B*_*trial*_, *D*_*trial*_ values were 5 Å^2^ and 0.25 Å, respectively. The values found by *Phenix* (Afonine *et al*., 2018) and *Relion* (Zivanov *et al*., 2018) were considered as the control values *B*_*atom,n*_, *D*_*atom,n*_.

Fig. 12 shows that for the main chain atoms and the most of side chains atoms the estimated resolution *D*_*vic*_ is close to the values *D*_*atom*_ obtained with *Relion*. The exception are the side chains for which the density in experimental map is poor (Lys 43, Arg 48, Gln 65, Glu 67, Lys 71).

**Fig. 12.**
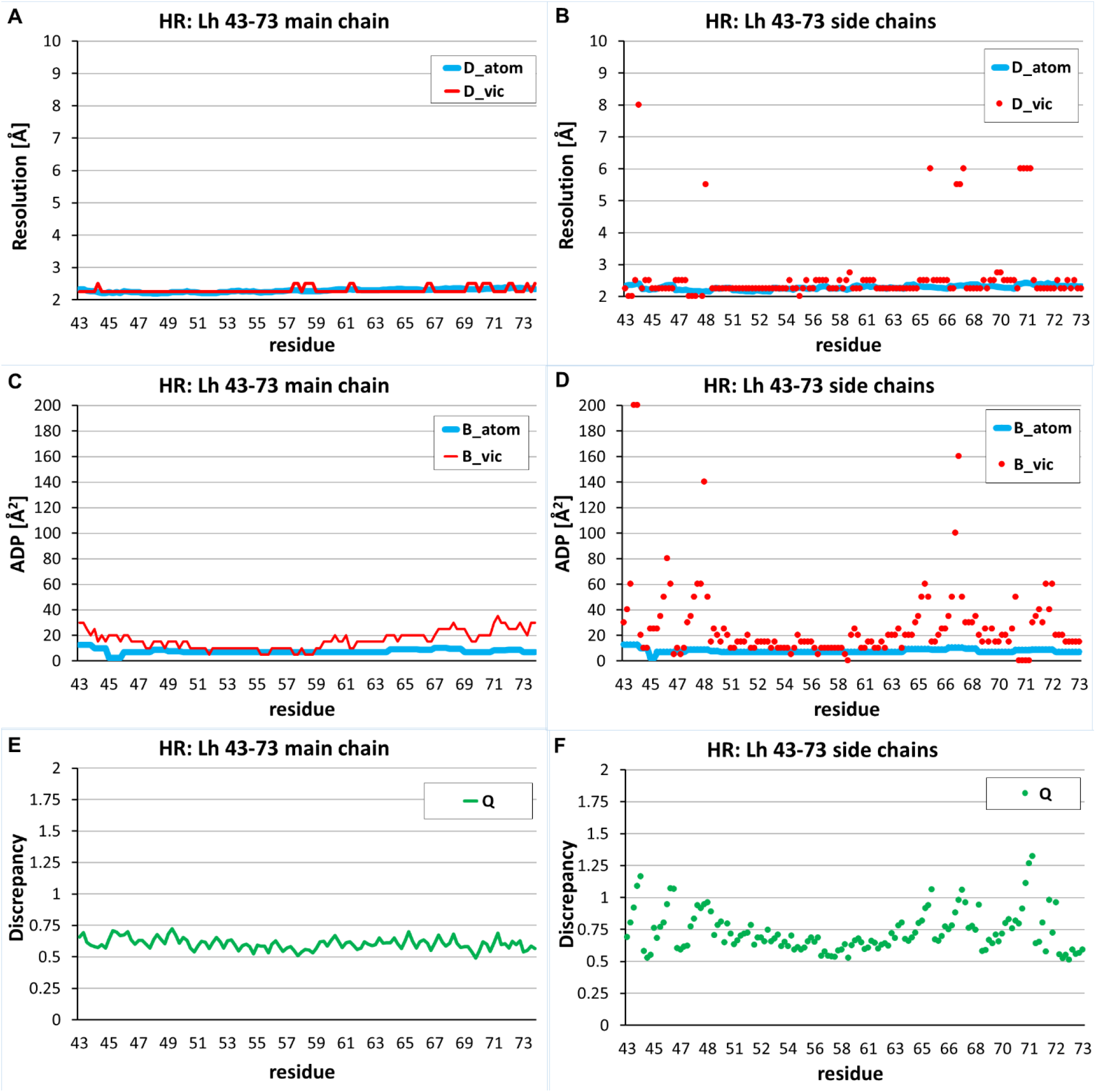
Results of heterogeneity analysis of the human ribosome cryo-EM map performed for Lys43-Tyr73 fragment (in chain Lh) shown for the main chain (left) and the side chain (right) atoms.

The found *B*_*vic*_ values for the main chain and most of side chain atoms vary around 20 Å^2^, the value usual for crystallographic atomic models obtained at 2 Å resolution. They are different from excessively low values *B*_*atom*_ defined by *Phenix*. For the six side chains with poorly defined density in the experimental map, the same as above plus Lys 46, the found *B*_*vic*_ values are much larger. The discrepancy (9) values corresponding to the optimal choice of *B*_*vic*_, *D*_*vic*_ reflect this tendency revealing larger Q-values for the atoms poorly defined in the map.

Fig. 13 shows a fragment of the experimental map (Fig. 13, A), and the same fragment of the map calculated with the atomic parameter values provided by *Relion* and *Phenix* (Fig. 13, B), and of the map calculated with the *B*_*vic*_, *D*_*vic*_ values found by the method suggested in this work (Fig. 13, C). The map calculated with the *Relion* and *Phenix* parameters does not reflect the actual quality of experimental map, *e.g*., for residues Glu 65, Glu 67, Lys 71. Instead, the parameters values *B*_*vic*_, *D*_*vic*_ obtained with the approach developed here allow reproducing the experimental map much better (Figs. 13 C, D).

**Fig. 13.**
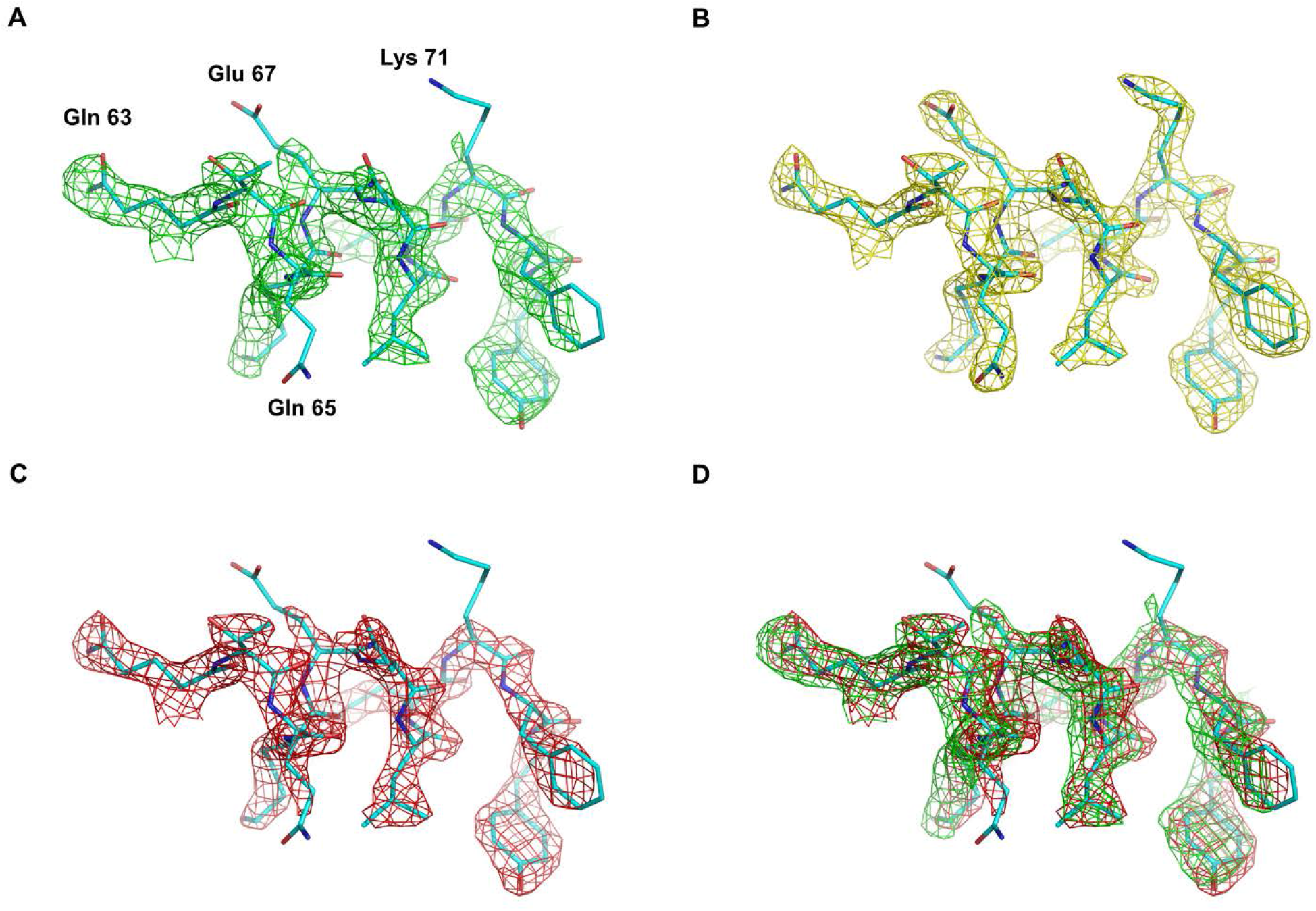
Maps showing the Lys43-Tyr73 fragment (chain Lh) superposed with the atomic model. A) The experimental cryo-EM map. B) The map calculated with the *B*_*atom*_, *D*_*atom*_ parameter values defined with *Relion* and *Phenix*. C) The map calculated with the *B*_*vic*_, *D*_*vic*_ parameter values, defined by minimization of discrepancy (9). D) Superposition of maps from A and C. The cut-off value in each map was chosen to display the equal specific volume of the selected regions (0.15 Å^3^Da^-1^).

It is worthy of noting that the atomic model used in this test is not the ideal model corresponding to the experimental map. Eventual inaccuracies in the atomic positions can be revealed by comparison of the experimental map with the image calculated as (6-7) with the parameters *B*_*vic*_, *D*_*vic*_ defined by minimization of (9). Fig.14 gives an example of such comparison suggesting the developed approach to be a component of future real-space refinement procedures allowing to improve atomic models further.

**Fig. 14.**
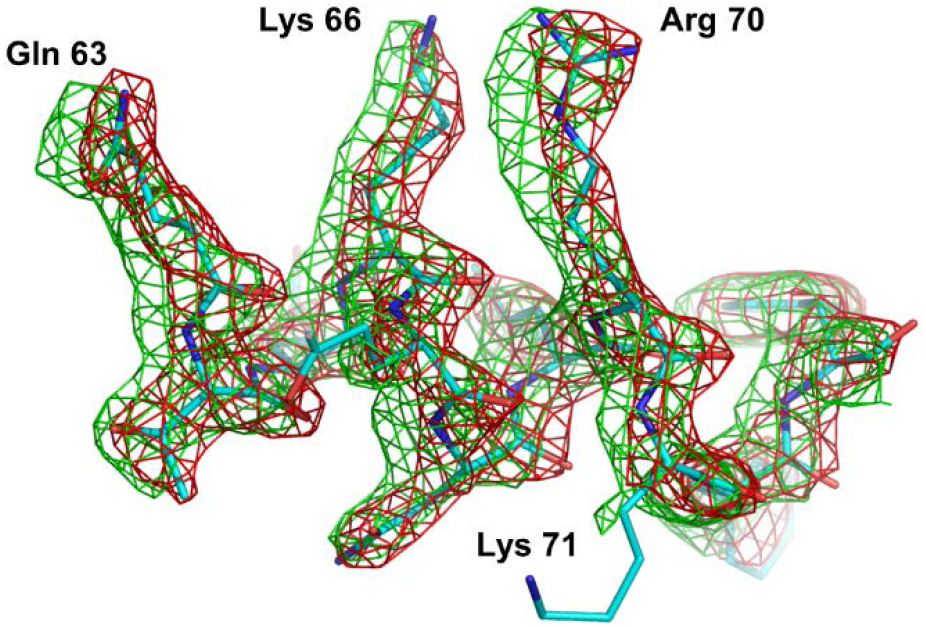
A fragment of the atomic model superposed with the experimental cryo-EM map (green) and with the model map (red) calculated with the current values of the atomic parameters. The revealed difference suggests to correct the position of some atoms. The cut-off in both maps was chosen to display the equal specific volume of the selected regions (0.15 Å^3^Da^-1^).

## 4. Discussion

Experimental maps in structural biology usually differ from the respective theoretical distributions due to various reasons. In this paper we address to two important reasons, namely a positional disorder of atomic centers and a limited resolution with which images of atoms appear in experimental maps.

These two characteristics may be heterogeneous varying from one region to another. Eventually they may be attributed individually for every atom of the object as atomic displacement parameter *B*_*atom*_ and atomic image resolution *D*_*atom*_. Recently we have suggested a method to describe atomic images of a limited resolution by shell-functions, that allows one to calculate heterogeneous density maps (Urzhumtsev and Lunin, 2022a) as sums of images of isolated atoms expressed analytically in terms of heterogeneity parameters.

In this article, we report the results of our attempts to develop an inverse procedure, *i.e*., to annotate heterogeneity of an experimental density maps in terms of point heterogeneity parameters of all atoms. In a future, this could be done by a multidimensional search in the space of all atomic heterogeneity parameters, minimizing the discrepancy between the calculated and experimental maps. At the moment, we have proposed a simplified approach that allows one to obtain estimates of these parameters locally, processing atom by atom. The problem in localization of the search is that in every point in space the value of the density is a sum of several atomic contributions, mixing up parameters of neighboring atoms in the discrepancy function. To overcome this problem, we introduce the concept of the local map heterogeneity parameters *B*_*vic*_, *D*_*vic*_ assuming temporary that in a vicinity of a given point all atoms contributing to the map have the same point heterogeneity parameters. The values *B*_*vic*_, *D*_*vic*_ may be assumed a ‘consensus’ values of individual parameters of neighboring atoms. These map heterogeneity values may be calculated locally for every point in space and produce some kind of annotation of the heterogeneity of the given density map. One should distinguish between then map heterogeneity parameters *B*_*vic*_, *D*_*vic*_, calculated from the map values around the center of an atom, and the point parameters *B*_*atom*_, *D*_*atom*_ with which this atom contributes to the map. Therefore, the first goal of our study was to verify whether the calculated *B*_*vic*_, *D*_*vic*_ values may serve as estimates for the point heterogeneity parameters. Our test, conducted for simulated maps, where the only present types of distortions were positional disorder and the loss of resolution (Sec. 3.1, 3.2), gave the positive answer for a vast majority of atoms. The ambiguity of the definition of parameters was manifested only for highly disordered low-resolution parts of the maps (Fig. 5).

The experimental methods of structural biology such as X-ray and neutron crystallography and single particle cryo-EM, report neither *B*_*vic*_, *D*_*vic*_ nor *B*_*atom*_, *D*_*atom*_ directly. Moreover, experimental maps are distorted also by the reasons other than the positional disorder and loss of resolution.

Therefore, the second goal of tests, performed this time with experimental maps (Sec. 3.3, 3.4), was to find to what extents *B*_*vic*_, *D*_*vic*_ calculated with the proposed method reproduce the values of atomic displacement parameter *B*_*XR*_ obtained by crystallographic refinement programs or *D*_*EM*_ resolution estimated by the Fourier Shell Correlation function in cryo-EM.

For experimental X-ray maps, the difference between *B*_*vic*_ values and *B*_*XR*_, extracted from PDB, did not exceed those in the tests with simulated maps (Fig. 10 A and 11 A *vs* Fig. 4A). This suggests both that the values *B*_*vic*_ are a reasonable approximation to the crystallographic displacement parameters *B*_*XR*_, and that the additional distortions inherent in the map do not significantly affect the accuracy of this approximation. The *D*_*vic*_ value in different points coincided with the formal resolution of Fourier syntheses for most atoms, or was only slightly different from it (Fig. 11 B). The exceptions were poor defined regions in the map, where the variation of resolution improved the correspondence between the calculate and experimental maps. Surprisingly, in all our tests, the resolution was estimated more accurately than the ADP values (Fig. 6). We hypothesize that this may be explained by the fact that a variation of the resolution displaces the positions of the ripples, which may be a stronger information than the degree of blurring of the density peaks.

For well-defined regions in the analyzed cryo-EM map, the found *D*_*vic*_ values were close to the *D*_*EM*_ values annotated by *Relion* program (Fig. 12 A, B). Nevertheless, for some side chains poor defined in the map *D*_*vic*_ differed significantly from the *D*_*EM*_ values, suggesting much lower resolution (Fig. 12 B). The calculation of fragments of the model map with {*B*_*EM*_, *D*_*EM*_} replaced by {*B*_*vic*_, *D*_*vic*_} showed that using the new values of the parameters allows one to reproduce the experimental map better (Fig. 13). Concerning the displacement parameters, our approach has resulted in usual-scale values *B*_*vic*_ which are larger than extremely small values *B*_*EM*_ annotated after previous refinement against experimental cryo-EM map (Fig. 12 C, D). There are two possible explanations. Firstly, EM-maps are subjected to additional processing, for example, sharpening, and to describe such maps, atomic displacement parameters should be also reduced. Secondly, the refinement against cryo-EM maps is not yet a well-established procedure requiring its further development.

The proposed approach to estimate the heterogeneity of maps uses available values of the atomic coordinates. If these coordinates contain errors, then a map corresponding to the model calculated with the found heterogeneity parameters can reveal errors in the location of atoms (Fig. 14). Inversely, incorrect atomic position may lead to errors in estimated *B*_*vic*_, *D*_*vic*_ values. A joint refinement of all parameters may be an option in future.

A significant problem when working with experimental maps is scaling, *i.e*., bringing the map calculated from the model to the experimental scale by a linear transformation *ρ*^*new*^ = κ(*ρ*^*model*^ − *ρ*^0^). The proposed approach to search for *B*_*vic*_, *D*_*vic*_ has a local character, *i.e*., each calculation is done independently in a small vicinity of a point. An attempt to determine the optimal scaling coefficients also locally, based on the analysis of the map values only inside this vicinity, can lead to a sharp variation of the values of scaling parameters between neighboring vicinities (Fig. 7A, Fig. 8 C, D, Fig. 10A). This in turn may results in errors in the determined heterogeneity parameters. The behavior of parameter κ changes drastically if the parameter *ρ*^0^ is fixed for all vicinities (Fig. 7) being estimated as

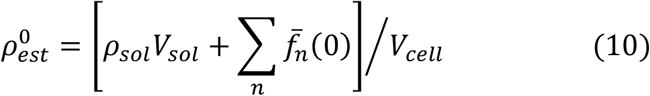

Here 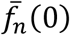 are atomic scattering factors (2) or (4) in the origin, *V*_*cell*_ is the unit cell volume, the sum is calculated over all atoms in the unit cell and the complementary term is designed to consider the bulk solvent contribution, when exists, with *ρ*_*sol*,_*V*_*sol*_ being the solvent density and the volume of the solvent region.

The results obtained in the reported work prove the feasibility of the solution of the inverse problem of the decomposition of density maps, *i.e*., a possibility to recover the values of the parameters of two principal phenomena which affect the molecular images, namely the local resolution and the local positional uncertainties. For well-ordered regions, these parameters may be included as variable to be adjusted by refinement programs. For poorly-ordered regions various acceptable combinations of the respective parameters lead to similar density maps and any of these combinations can be taken. This aspect of the procedure, as well as modeling of more tiny effects such as an anisotropic disorder, require a further analysis which is in progress.

## 5. Conclusions

The proposed approach allows to annotate the heterogeneous imperfections of experimental maps obtained in structural biology. Within this approach, the heterogeneity is expressed in terms of variation, from one point to another, of the local atomic displacement and local resolution parameters. The developed real space procedure, based on the shell approximation of atomic images, allows appropriate estimates of these parameters, and reproduces properly the resolution value when applied to maps of a homogeneous resolution. For well-ordered parts of the structure, the estimates of the atomic displacement parameters determined by the proposed method are close to those found by reciprocal space refinement. For less ordered parts, which are poorly represented in an experimental map, a simultaneous correction of displacement and resolution parameters presents an alternative description of heterogeneity and allows one to get a better local correspondence of the calculated and the observed maps.

## CRediT authorship contribution statement

Vladimir Y. Lunin: Conceptualization, Algorithms, Discussions, Writing. Natalia L. Lunina: Calculations, Discussions. Alexandre G. Urzhumtsev: Conceptualization, Algorithms, Discussions, Writing.

## Acknowledgements

We thank L.M. Urzhumtseva for help with computing, tests, and program development. AU acknowledges Instruct-ERIC and the French Infrastructure for Integrated Structural Biology FRISBI [ANR-10-INBS-05].

This research did not receive any specific grant from funding agencies in the public, commercial, or not-for-profit sectors.

